# Discovery and synthesis of hydroxy-L-proline based blockers of the neutral amino acid transporters SLC1A4 (ASCT1) and SLC1A5 (ASCT2)

**DOI:** 10.1101/2021.12.14.470456

**Authors:** Brent R. Lyda, Gregory P. Leary, Jill Farnsworth, Derek Silvius, Benjamin Seaver, C. Sean Esslinger, Nicholas R. Natale, Michael P. Kavanaugh

## Abstract

The conformationally restricted heterocycle hydroxy-L-proline is a versatile scaffold for the synthesis of diverse multi-functionalized pyrrolidines for probing the ligand binding sites of biological targets. With the goal to develop new inhibitors of the widely expressed amino acid transporters SLC1A4 and SLC1A5 (also known as ASCT1 and ASCT2), we synthesized and functionally screened a series of hydroxy-L-proline derivatives or ‘prolinols’ using electrophysiological and radio-labeled uptake assays on amino acid transporters from the SLC1, SLC7, and SLC38 solute carrier families. We identified a number of synthetic prolinols that act as selective high-affinity inhibitors of the SLC1 functional subfamily comprising the neutral amino acid transporters SLC1A4 and SLC1A5. The active and inactive prolinols were computationally docked into a threaded homology model and analyzed with respect to predicted molecular orientation and observed pharmacological activity. The series of hydroxy-L-proline derivatives identified here represents a new class of potential agents to pharmacologically modulate SLC1A4 and SLC1A5, amino acid exchangers that play important roles in a wide range of physiological and pathophysiological processes.

## Introductions

The amino-acid exchangers SLC1A4 (ASCT1)^1,2^ and SLC1A5 (ASCT2)^3,4^ are sodium dependent transmembrane proteins that mediate flux of neutral, generally polar L-amino acids. These proteins were first characterized and dnoted as ASC (Ala, Ser, Cys -selective) based upon their representative substrates alanine, serine, and cysteine underlying a Na-dependent transport across the plasma membrane of mammalian.^5^ Both transporters exhibit generally similar substrate selectivity but differ notably with respect to glutamine, a substrate of only SLC1A5.^1,3^ While transmembrane flux of amino acids mediated by these proteins is Na^+^-dependent, it occurs by an obligate exchange mechanism without stoichiometrically coupled current flow.^6^ SLC1A4 and SLC1A5 make up a functionally distinct subfamily within the SLC1 family, which contains seven paralogs in mammals. The remaining five family members are acidic amino acid (glutamate/aspartate) transporters (SLC1A1-3 and SLC1A6-7; also known as excitatory amino acid transporters EAAT1-5).^7^ In contrast to the neutral amino acid exchangers SLC1A4/5, the other SLC1 paralogs are electrogenic and mediate coupled flux of amino acids, with co-transport of three Na^+^ ions and one proton and countertransport of one potassium ion.^8^ Despite the difference in coupled ion stoichiometry, all members of the SLC1 family mediate a chloride conductance that is not thermodynamically coupled to substrate flux. This anion current is distinct from the stoichiometrically coupled currents in the acidic amino acid transporter SLC1 subfamily and is the sole current measured in the neutral amino acid SLC1 exchanger subfamily.^6,9^

There are significant gaps towards our understanding of the physiological roles for the SLC1A4/5 transporters. SLC1A5 is widely expressed in tissues outside of brain and plays a role in the homeostasis of the centrally important amino acid glutamine.^3,4^ SLC1A5 has also been a focus of investigation in cancer pathophysiology because of its increased expression in several types of tumors.^10–12^ While SLC1A4 is also widely expressed in mammalian tissues, in contrast to SLC1A5, it is most highly expressed in brain.^1,2^ Mutations in the corresponding human gene have been associated with neurological disorders including cognitive and developmental impairment.^13–15^ In addition to transport of L-amino acids, SLC1A4 has recently been shown to mediate transport of D-serine,^16^ a co-agonist of the NMDA receptor, which has roles in synaptic transmission, memory, and neural development.

Developing pharmacological tools to assess the physiological function of SLC1A4 and SLC1A5 transporters is an important aim. There has been limited progress thus far in developing inhibitors and probes for these transporters with high potency and selectivity. SLC1A4/5 transporters exhibit affinities for their preferred neutral amino acid substrates in the range of 20– 200 μM.^1,3^ Non-transported SLC1A5 competitive inhibitors based on L-*γ*-glutamyl-*p*-nitroanilide or benzyl-serine and -cysteine derivatives exhibit K_i_ values ranging from 20-800 μM.^17–19^ More recent work has built off a proline scaffold with improvements in affinity to the lower micro molar range.^20,21^

We sought to generate new SLC1A4/5 ligands with high affinity and selectivity using a conformationally constrained scaffold based on hydroxy-L-proline. The neutral amino acid L-proline does not elicit detectable chloride currents at concentrations up to 1 mM in oocytes expressing SLC1A4.^1^ Nevertheless, application of extracellular 300 μM L-proline is capable of inducing heteroexchange release of intracellular radiolabeled ASCT substrates, indicating that it is a transportable ligand/substrate.^6^ Pinilla-Tenas and colleagues showed that a pyrrolidine analog of L-proline, the 4-positioned alcohol *trans*-4-hydroxy-L-proline (**2**), increased the apparent affinity for SLC1A4 by approximately 20-fold over L-proline.^22^ From a synthetic and computational perspective, pyrrolidine heterocycles are particularly useful for designing amino acids containing multiple stereocenters with unique spatial and conformational arrangements. We now report the synthesis, computational docking, and functional characterization of a novel series of hydroxy-L-proline derivatives (prolinols). Using electrophysiological approaches, we observe that several of these prolinols demonstrate pharmacologic activity targeting mouse or human isoforms of SLC1A4/5 transporters with affinities in the low micromolar and nanomolar range.

## Methods

### Chemicals and Reagents

L-[^3^H]-Alanine and L-[^3^H]-glutamine were purchased from Dupont NEN (Boston, MA). *trans*-4-Hydroxy-L-proline (4-HP, **2**), *trans*-3-hydroxy-L-proline (3-HP), and benzyl alcohols were obtained from Acros Organics (Morris Plains, NJ). LC-MS grade acetonitrile for HPLC was obtained from EMD Chemicals USA. All other chemical reagents were purchased from Acros Organics (Morris Plains, NJ). All ACS reagent grade EMD brand solvents were purchases from VWR International (Radnor, PA). Separation of regioisomers and isolation of final products were performed on a HPLC C18, 250mm x 21.2mm, 10μ column using a Waters 486 Tunable Detector, and a Milton Roy pump. Full experimental details are reported in the experimental section and supplemental sections.

### Expression and functional testing of amino acid transporters expressed in Xenopus laevis oocytes

The molecular pharmacology of amino acid transport systems SLC1A1-5 (EAAT3, EAAT2, EAAT1, SLC1A4 (ASCT1), murine SLC1A5 (ASCT2), human SLC38A1 (ATA1), SLC38A2 (ATA2), SLC38A4 (ATA3), and SLC7A10+SLC3A2 (asc-1+4f2) were characterized by heterologous expression in *Xenopus laevis* oocytes. Radiolabeled uptake and electrophysiology measurements were made as previously described.^6^ Briefly, oocytes were injected with approximately 50 ng of cRNA transcribed from cDNA. For radiolabeled uptake competition experiments, oocytes injected with transporter cRNA were incubated over 10 minutes in frog ND96 buffer with indicated concentrations of [^3^H]L-alanine or [^3^H]L-glutamine (American Radiolabeled Chemicals; 40-60 Ci mmol^−1^) with or without competing ligands. Uptake was halted by washing 3 times with 4°C ND96 buffer. Oocytes were then lysed in 0.1% Triton X-100, and radioactivity was counted via liquid scintillation. Dose response curves were generated by fitting fractional inhibition (I) to the expression I = I_max_ * [inhibitor]/([inhibitor]+IC_50_). Inhibition of radioligand uptake at single concentrations was performed using 1mM of test compound or control ligand to determine percent inhibition. For radiolabeled heteroexchange assays, oocytes were incubated for 20 minutes with radiolabeled 100 μM [^3^H]L-alanine (4 μCi / mL) in frog ringer!s solution. Ligand substrates or non-substrate inhibitors, as indicated, were then assayed for the capacity to induce exchange/release of accumulated radiolabeled alanine over 20 minutes by transferring 5 oocytes/well to 500 μL frog Ringer!s solution containing 1mM of test ligand.

Two-electrode voltage clamp (TEVC) transporter currents were recorded with a GeneClamp 500 amplifier and Digidata analog-digital computer interface (Molecular Devices). Electrodes (0.2–1.0 MΩ) were filled with 3M KCl and the recording chamber was grounded with an agar bridge. Data were acquired with Axograph and LabChart and analyzed with Kaleidagraph software (Synergy). Uncoupled anion currents associated with the transport of substrate and block of leak conductances mediated by ASCT1 and mASCT2 were recorded in the presence of Ringer with 50% substitution of NaCl by NaSCN to amplify the non-transported anion conductance.^6^

### Computational Docking

SLC1A5 or SLC1A4 primary sequence (GenBank accession number D85044 or L14595 respectively), was aligned with the Protein Data Bank (PDB) sequences for the archaeal homologue Glt_Ph_ co-crystallized with non-substrate inhibitor L-*threo*-β-benzyloxyaspartate TBOA, 2NWW.pdb.^23^ The ASCT2 homology model was constructed by threading the aligned sequence along 2NWW coordinates using T-Coffee–Fasta alignment via SCWRL – server (http://www1.jcsg.org/scripts/prod/scwrl/serve.cgi. The resulting homology model was optimized through local energy minimizations of regions with high steric and electrostatic interference using the AMBER7 force field in SYBYL (Suite, 2010). Energy minimized prolinols using SYBYL [new reference] were docked using GOLD [reference] into the homology model. Each of the top ten poses for each of three separate docking experiments was evaluated for their capacity to dock with ASCT2 residues C479 and D476 or in ASCT1 residues T459 and D456. The top GOLDScore poses and lowest predicted –ΔG (kJ/mol) values were merged into the homology model and visualized via PyMOL Molecular Graphics-System-Version 1.3, SchrödingerLLC.

## Results

### Hydroxy Proline Scaffold

*Trans*-3-hydroxy-L-proline (3-HP) and *trans*-4-hydroxy-L-proline (4-HP) were evaluated as substrates of SLC1A4 and murine SLC1A5 (mSLC1A5) with two-electrode voltage clamp (TEVC) measuring substrate induced SCN^-^ anion currents, a technique found to be reliable for screening and characterizing transport kinetics of the SLC1A4/5 transporters.^6,9^ Application of 3-HP or 4-HP to oocytes expressing SLC1A4 or mSLC1A5 revealed anion currents that were concentration dependent and saturable. These dose response curves were normalized to the maximal response induced by the principal substrate, L-Ala or L-Gln, respectively (Figure 1). The dose responses were fit by the Michaelis-Menten equation (Figure 1A_ SLC1A4: 3-HP Km = 14.0+/- 1.8μM; 4-HP Km = 17.1 +/- 0.8μM; L-Ala Km 40.0 +/- 1.8μM or Figure 1B_ mSLC1A5: 3-HP Km = 28.6 +/- 6.4μM; 4-HP Km = 76.3 +/- 11.0μM, L-Gln Km = 47.4 +/- 4.6μM). For both molecules, there was a higher apparent affinity for SLC1A4 compared with mSLC1A5. Radiolabeled uptake of [H^3^]L-Gln was dose dependently blocked by 3-HP and 4-HP, confirming the affinity for mSLC1A5. Cross reactivity was tested by applying 4-HP at 1 mM to glutamate transporters SLC1A3 or SLC1A1. No TEVC current response or block was measured suggesting a selectivity within this SLC1A gene family (data not shown). 4-HP did induce currents in the neutral amino acid transporter SLC38A2 but with a 100 fold lower affinity (SLC38A2_4-HP Km = 3.1mM_data not shown).

**Figure 1.**
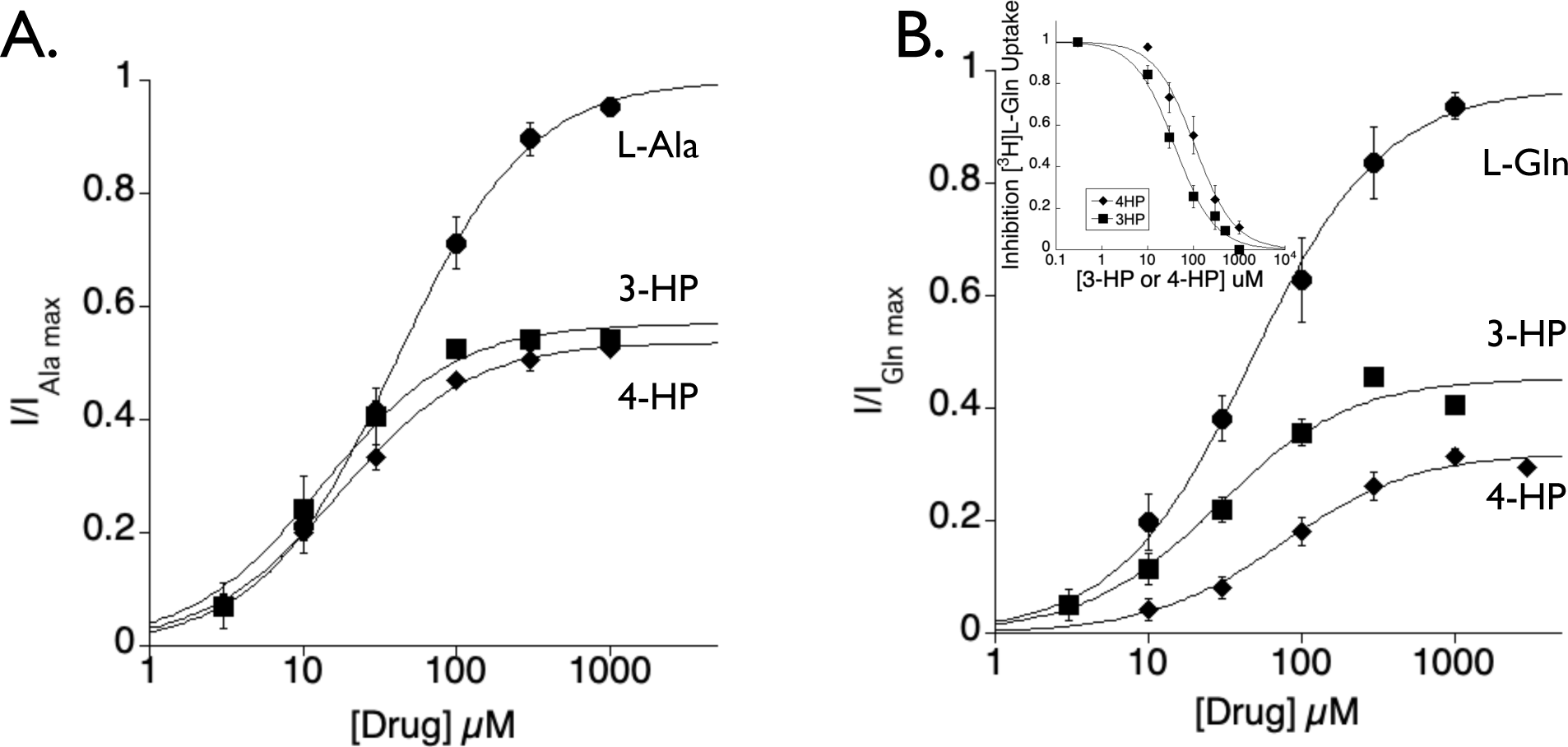
The scaffolds 3-HP and 4-HP dose dependently activate SLC1A4 and mSLC1A5 transporter currents. **A**. SLC1A4 dose response curves for substrate activated anion currents using two-electrode voltage clamp of *Xenopus lavies* oocytes (−30mV). Currents are normalized to the maximum current estimated for alanine from the Michaelis-Menten equation fit (Ala Km = 40.3+/-1.8; 3-HP Km = 13.5 +/-1.8; 4-HP Km = 17.1 +/-0.8). **B**. mSLC1A5 dose response curve for substrate activated anion currents using two-electrode voltage clamp (−30mV). Currents are normalized to the maximum current estimated for glutamine from the Michaelis-Menten equation fit (Gln Km = 47.4+/-4.6; 3-HP Km = 28.6 +/-6.4; 4-HP Km = 76.3 +/-11.0). Extracellular solutions contained 50 mM NaSCN. *Inset B:* [^3^H]L-Gln uptake in mSLC1A5 is blocked dose dependently by both 3-HP (IC50 = 39uM) and 4-HP (IC50 = 108uM).

### Inhibitor Design

#### Prolinol Derivatives

Hydroxy-proline derivatives are appealing for ligand design because of the conformationally restricted pyrrolidine ring (Figure 2). A series of diverse analogs were designed with an array of various stereo- and regiochemical configurations.

**Figure 2.**
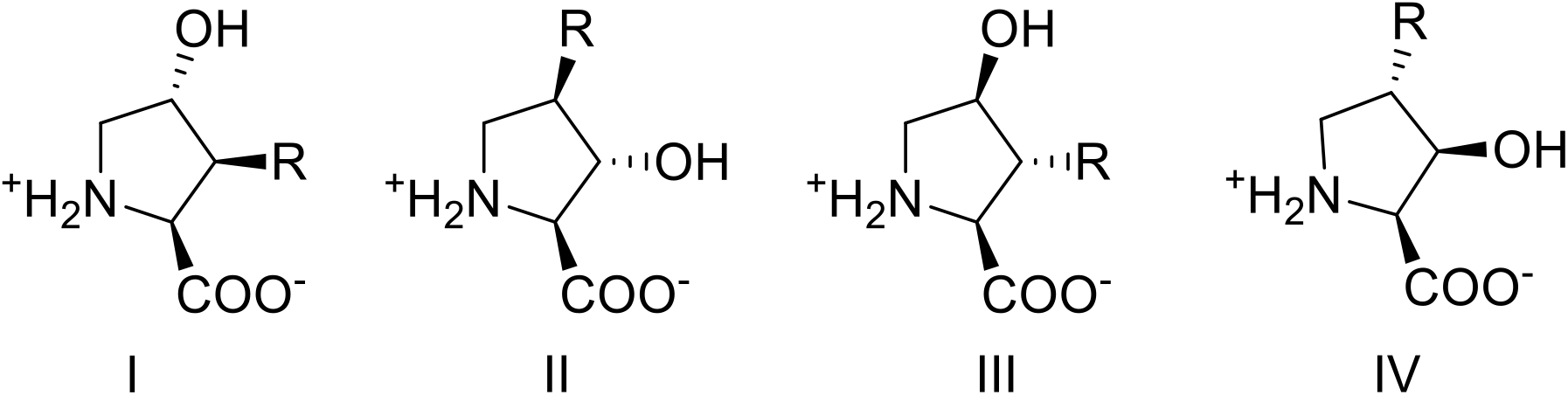
Multi- functionalized hydroxy prolinol targets to inhibit SLC1A4 and mSLC1A5. R = any group.

A series of multi-functionalized pyrrolidines were synthesized from protected *cis* or *trans*-3,4-epoxy-L-prolines including regioisomers of methyl, phenyl or phenol-ether substituted hydroxy-L-prolines, compounds **10**[**a-d**], **12a & 12b**; benzyl or alkyl ether substituted prolinols, compounds **14**[**a-f**], **17**[**a-c**]; and finally, derivatives of N-protected 4-keto-proline, *cis*-4-methyl-*trans*-4-hydroxy-L-proline (**26**) and *cis*-4-hydroxy-*trans*-4-methyl-L-proline (**27**).

The stereochemistry and regiochemistry of these 3,4-substituted prolinol products were established from the fully protected (N-Cbz & benzyl ester) 3,4-epoxy-prolines from which they were derived. The *cis* and *trans* epoxide isomers are easily isolatable allowing stereochemistry to be determined by 2D-NOE. Our stereo-chemical assignments of both isomers are consistent to that previously reported.^24^ Stereochemistry of geminal substituted 4-methyl-4-hydroxy-substituted prolinols was deduced directly as the final products by 2D-NOE. Regio chemical assignments of final product hydroxy-proline derivatives were inferred by a combination of 2D gradient correlation spectroscopy (gCOSY) for all final products in addition to 2D-NOE (for **14a, 14b** and **17a** only). Supplemental Fig. 1 illustrates how the proton assignments and regiochemistry were made from a gCOSY experiment of example prolinol *trans*-3-hydroxy-*cis*-4-isopropoxy-L-proline **14b**. Refer to the experimental procedures section for full experimental details, procedures, and synthetic schemes.

**Scheme 1.**
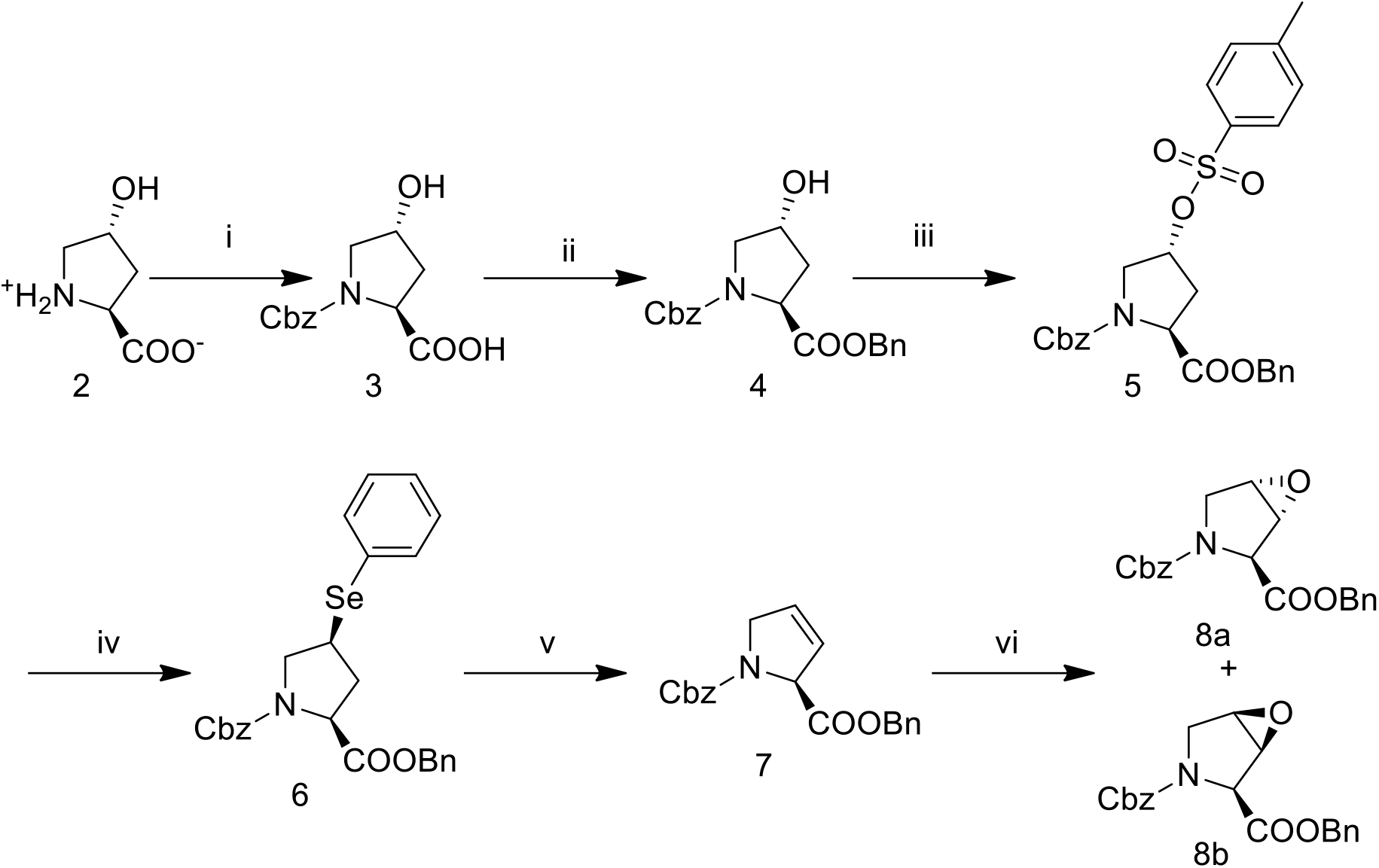
Synthesis of N-Cbz-*trans*- and *cis*-3,4-epoxy-proline-benzyl ester **8a & 8b**. (i) benzylchloroformate, H_2_O, NaOH, rt; (ii) benzylbromide, Na_2_CO_3_, NaI, rt; (iii) p-toluenesulfonyl-Cl, pyridine, 0°C; (iv) tBuOH, phenylselenide, reflux; (v) CH_2_Cl_2_, H_2_O_2_, 0°C-RT; (vi) mCPBA, CH_2_Cl_2_, reflux

### Inhibitor Chemistry

#### Preparation of cis-4-hydroxy-L-proline

Analog *cis*-4-hydroxy-L-proline (1) was synthesized from *trans*-4-hydroxy-L-proline as previously reported. Briefly, the amine of *trans*-4-hydroxy-L-proline was protected using di-tert-butyl dicarbonate under standard conditions and aqueous workup. The isolated N-tBoc-*trans*-4-hydroxy-L-proline was then converted to the N-tBoc-*cis*-proline lactone by intramolecular Mitsunobu esterification using triphenyl phosphine (PPH_3_) and diethyl azodicarboxylate (DEAD) in THF. The N-tBoc-proline lactone was then converted to *cis*-4-hydroxy-proline in a one-step hydrolysis following heating in 2.0 M HCl, (2.2 eq) and isolation via ion exchange.

**Scheme 2.**
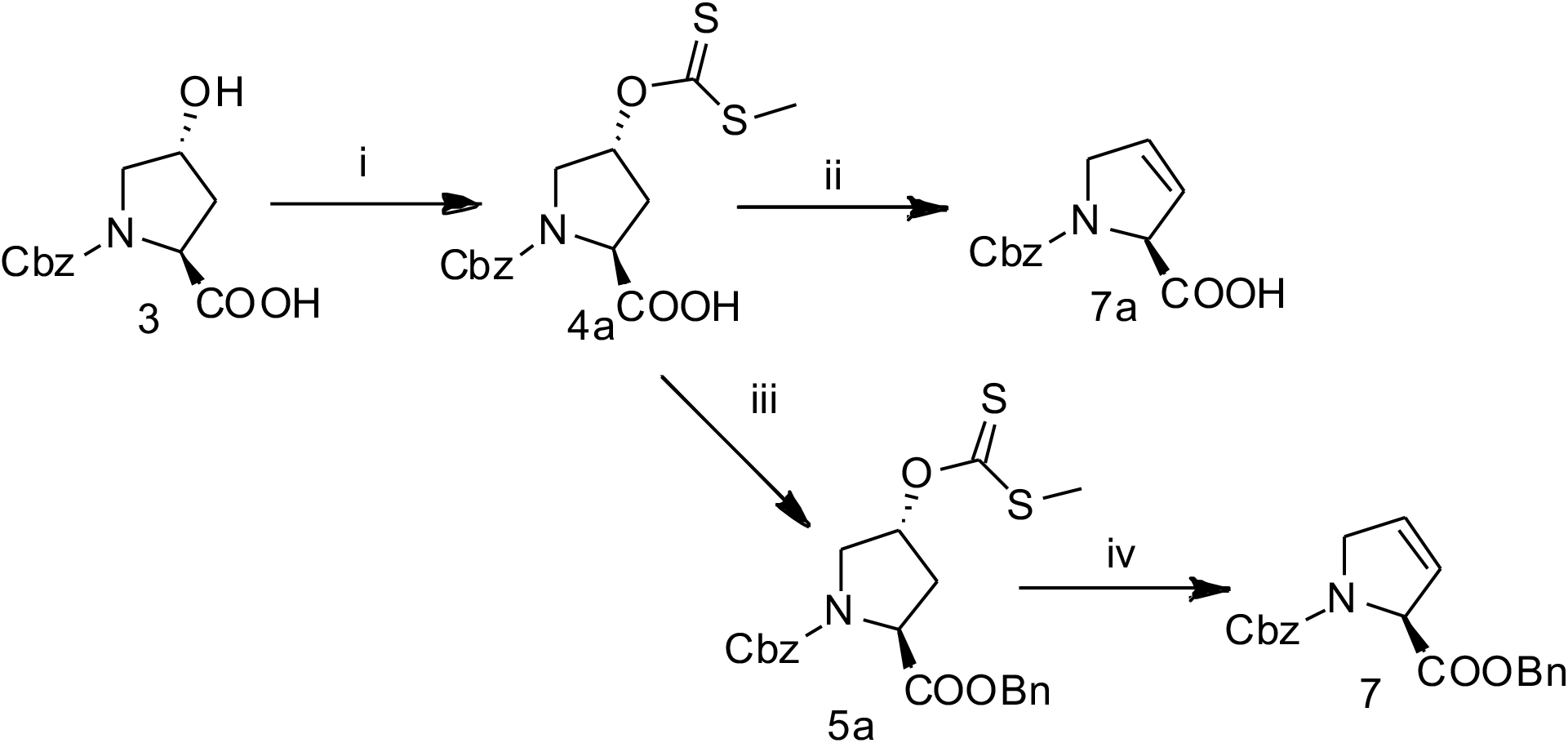
Alternative synthesis of 3,4-dehydroprolines **7a** and **7**. (i) NaH, THF, CS_2_, iodomethane, 22°C; (ii) H_2_O, NaHCO_3_, microwave 100°C; (iii) benzylbromide, Na_2_CO_3_, NaI, rt; (iv). H_2_O, Na_2_CO_3_, microwave 100°C.

#### N-Cbz-(cis- or trans)-3,4-epoxy-L-proline benzyl ester

To ultimately arrive at prolinol targets I-IV (Figure 2), we adapted a route to convert *trans*-4-hydroxy-L-proline (2) to the protected *trans*- and *cis*-3,4-epoxides (8a & 8b) illustrated in Scheme 1 with slight modification to that previously reported.^24^ Briefly, amine protection of *trans*-4-hydroxy-L-proline (2) was accomplished using benzylchloroformate in water at a pH ∼10. The carboxylic acid of N-Cbz-*trans*-4-hydroxy-L-proline (4) was then benzyl protected with benzylbromide, Na_2_CO_3_, NaI, in DMF. Intermediate 5 (N-Cbz-4-tosyl-L-proline benzyl ester) was obtained via treatment of the 4-alchol (4) with p-toluenesulfonyl chloride in pyridine at 4°C. Substitution of the tosylate (5) with reduced phenyl selenide, reflux in *tert*-butanol, provided the 4-phenylseleno-proline intermediate (6) and post isolation was treated with H_2_O_2_ to obtain the olefin, protected 3,4-dehydro-L-proline (7). Intermediate 7 was then directly converted to the protected *trans*- and *cis*-3,4-epoxy-prolines (8a & 8b; Scheme 4) or the benzylic ester was hydrolyzed to olefin intermediate 7a and converted to the epoxide (8c, Scheme 3) using *meta*-chloroperoxybenzoic acid (mCPBA). The fully protected *trans*- and *cis*-epoxides (8a or 8b) were efficiently isolated in modest yields, ∼40% up to this point within the synthetic scheme. Portions of the epoxides 8a or 8b were subsequently used in a BF_3_·Et_2_O initiated substitution (Scheme 4) whereas free acid epoxide intermediate 8c was coupled under basic conditions (Scheme 3).

**Scheme 3.**
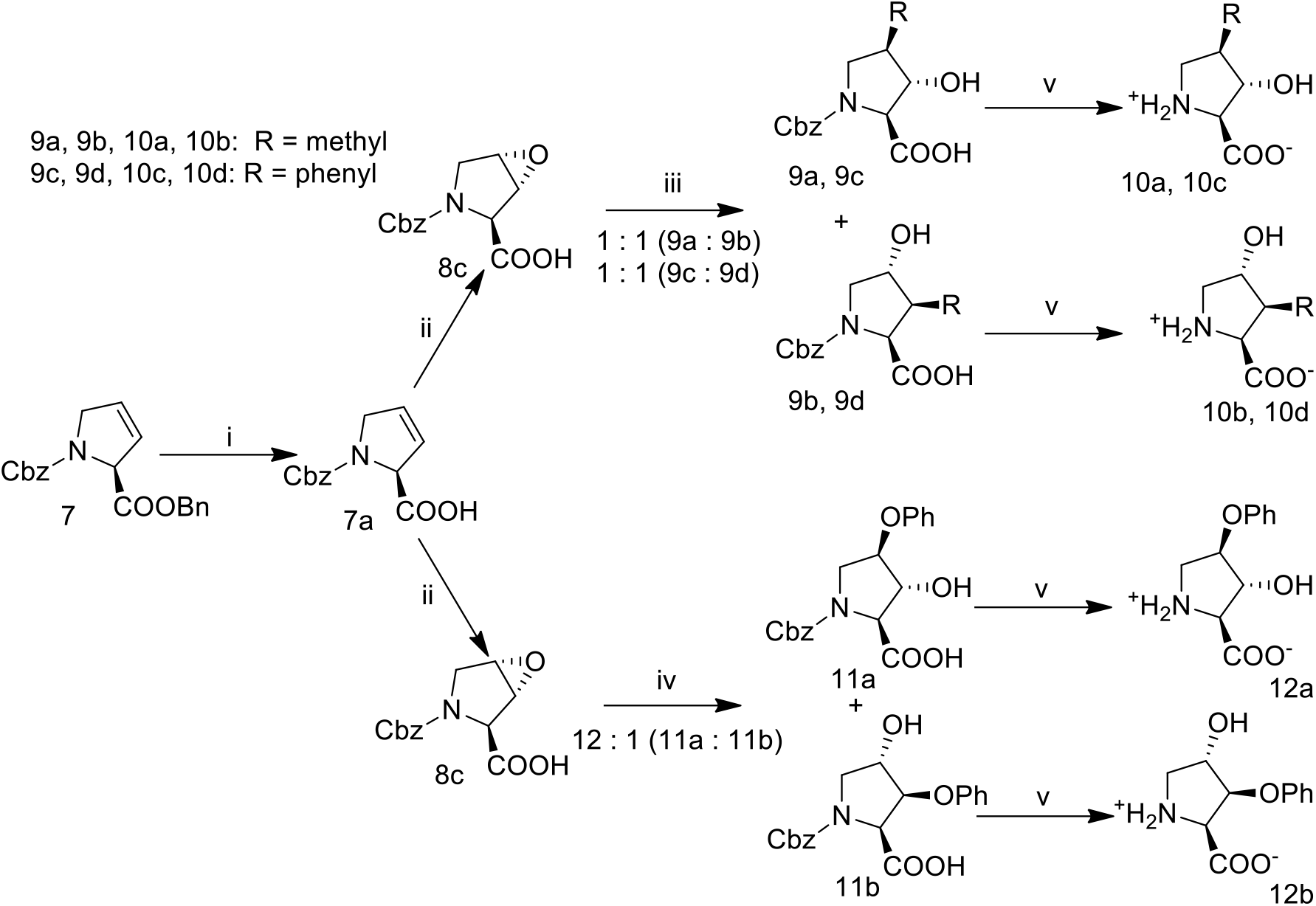
Synthesis of alkyl, aryl or phenol-ether prolinols. (i) KOH, THF : H_2_O, 0°C; (ii) mCPBA, CH_2_Cl_2_; (iii) 2 eq. R_2_Cu(CN)Li_2_, Et_2_O, -40°C under Ar (g); (iv) Sodium phenoxide in THF, 0°C - rt; (v) 10% Pd/C (10% weight. eq), 40 psi H_2_, MeOH, H_2_O, NH_4_OAc. 9a, 9b, 10a, 10b; R = methyl. 9c, 9d, 10c, 10d; R = phenyl.

#### Alternative synthesis of N-Cbz-3,4-dehydro-L-proline-benzyl ester 7 or N-Cbz-3,4-dehydro-L-proline 7a

An alternative route to produce 3,4-dehydro-L-prolines was explored through elimination of methyl xanthate ester intermediate. It was found that both free acid 3,4-dehydro-proline 7a or benzyl ester protected 3,4-dehydro-proline 7 could be generated via microwave assisted pyrolysis elimination of the methyl xanthate 4a or 5a, Scheme 2. Briefly, N-Cbz-*trans*-4-hydroxy-L-proline 3 was used in an esterification with CS_2_ and iodomethane using NaH in THF to produce N-Cbz-*trans*-4-methylxanthate-L-proline 4a. Carboxylic acid protection was achieved with benzyl bromide under standard conditions providing 5a, Scheme 2 (iii). Both methyl xanthate pyrrolidines 4a and 5a were successfully eliminated under microwave irradiation in an open / unsealed flask, with sodium carbonate and water to provide the 3,4-dehydroprolines 7 and 7a.

#### Methyl, phenyl or phenyl-ether prolinols

Alkyl, aryl and phenoxy substituted prolinols were generated via nucleophillic substitution. As precaution to avoid possible racemization of the α-chiral center during the addition step free acid pyrrolidine epoxide intermediated 8c (Scheme 3) was used. Treatment of the free acid olefin (7a) with mCPBA in DCM under reflux produced the *trans*-epoxide 8c in excellent yields with no detectable *cis*-epoxide. Epoxidation via dimethyldioxirane generated *in-situ* or using peracidic acid also afforded epoxide 8c, albeit, in low yield. The N-protected substituted prolinol intermediates 9a and 9b or 9c and 9d were obtained from the free acid epoxide pyrrolidine by transformation of the organolithium reagent to a higher order cyano-cuprate complexation between 2 equivalents of organolithium reagent and CuCN (Scheme 3; iii).^25,26^ The resulting prolinol products, 3 or 4 substituted prolinols, were recovered in yields of 90-95% and the regioisomers isolated via reverse phase HPLC in ratios of nearly 1:1.

Similarly, the N-protected phenoxy prolinols **11a & 11b** intermediates were obtained from the free acid *trans*-epoxide **8c** under basic conditions via addition of sodium phenoxide with regioisomeric ratios of 12:1, 3-hydroxy : 4-hydroxy respectively, with modest yields of ∼80%. Hydrogenolysis of the N-benzylcarbamate (Cbz) of protected prolinols **9a**-**d, 11a** and **11b** in AcOH and subsequent separation of the regioisomers via C18-reverse phase HPLC afforded the final prolinol products **10a**-**d, 12a** and **12b**.

**Scheme 4.**
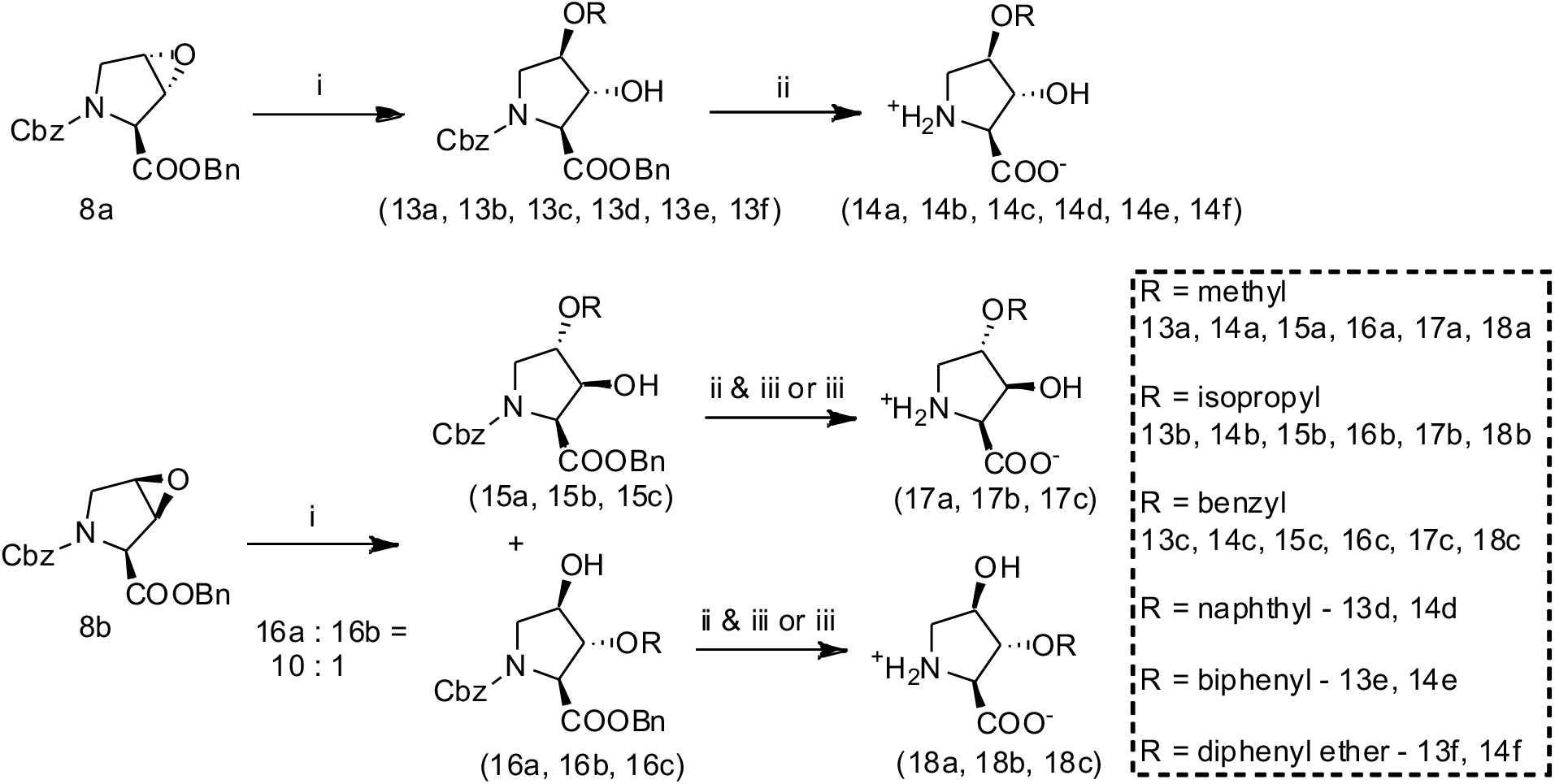
Synthesis of ether substituted prolinols. (i) BF_3_·Et_2_O, CH_2_Cl_2_, desired primary alcohol (HOMeR), 22°C; (ii) KOH, THF, H_2_O; (iii) 10% Pd/C (10% weight. eq.), H_2_, MeOH, H_2_O, ^+^NH_4-_OAc (0.5 eq). 13-18(a) R = methyl; 13-18(b) R = isopropyl; 13-18c R = benzyl; 13-14(d) R = naphthyl; 13-14(e) R = biphenyl; 13-14(f) R = diphenyl ether.

#### Ether substituted prolinols

Ether substituted prolinols were synthesized and explored as potential inhibitors of the SLC1A4/5 transporters. Assembly of this series was accomplished by a BF_3_•Et_2_O assisted nucleophillic ring opening of epoxide **8a** or **8b** with a primary alcohol and subsequent Pd catalyzed hydrogenolysis to produce *trans*-hydroxy derivatives **14(a-f)** or *cis*-hydroxy derivatives **17 & 18** (Scheme 4). BF_3_•Et_2_O initiated substitution of the epoxide provided superior yields compared to substitution initiated with *p*-toluenesulfonic acid which produced byproducts, including fortuitous addition of the tosylic acid later utilized to produce the product (2S,3R,4S)-3-hydroxy-4-phenoxypyrrolidin-2-carboxylate **12c**, see Supplemental figure 4. Ring opening of *trans*-epoxide **8a** under BF_3_•Et_2_O assisted substitution produced exclusively one regioisomer, the N-Cbz-*trans*-3-hydroxy-*cis*-4-ether-L-proline benzyl ester **13**. Whereas the ring opening of the *cis*-epoxide **8b** produced both regioisomer intermediates **15 & 16** in ratios, 8:1 (approximated by ^1^H NMR), greatly favoring the 3-positioned alcohol intermediates **15(a-c)**, consistent with azide or HCl ring opening of these epoxides.^24,27^ Separation of these regioisomers was carried out by reverse phase HPLC following hydrolysis of the benzyl ester with KOH or just after hydrogenolysis. Of note, though not explored here, we’ve observed that *cis*-3-alkyl-ether-*trans*-4-hydroxy-L-proline regioisomers of **14** may also be obtained from the free acid *trans*-3,4-epoxy-proline derivative **8c** (Scheme 3) via BF_3_•Et_2_O initiated addition under similar conditions(results not shown). Benzyl carbamate deprotection of ether substituted prolinols **13, 15** or **16** was again accomplished via palladium catalyzed hydrogenolysis under H_2_ (g) affording final products **14, 17**, or **18**. Importantly, we found that selective hydrogenolysis could be achieved via addition of ammonium acetate (NH_4_OAc) as an amine-based inhibitor of debenzylation during palladium catalyzed hydrogenolysis.^28,29^

### Pharmacology

#### Hydroxy-prolinol derivatives inhibit SLC1A4/5

Inhibitors of SLC1A5 block an anion leak conductance.^9^ Our results with conformationally constrained prolinols confirmed this effect in both SLC1A4 and mSLC1A5. Figure 3 illustrates this block of the anion leak conductance which resulted in an inward as opposed to the outward current induced by substrates. As expected for conductance originating from the same channel, the inward and outward currents shared a reversal potential shown with the respective current/voltage relationship (Figure 3, inset). The current blocked by inhibitors was much more pronounced for mSLC1A5, revealing a larger leak conductance for this transporter. Each hydroxy-prolinol analog demonstrated a concentration dependent and saturable block of the outward leak current (Figure 3C & D). These dose response kinetics were independent of competing substrates, and thus provided a direct measurement of the inhibitor binding equilibrium constant (K_i_). Kinetic parameters for each compound are reported in Table 1.

**Figure 3.**
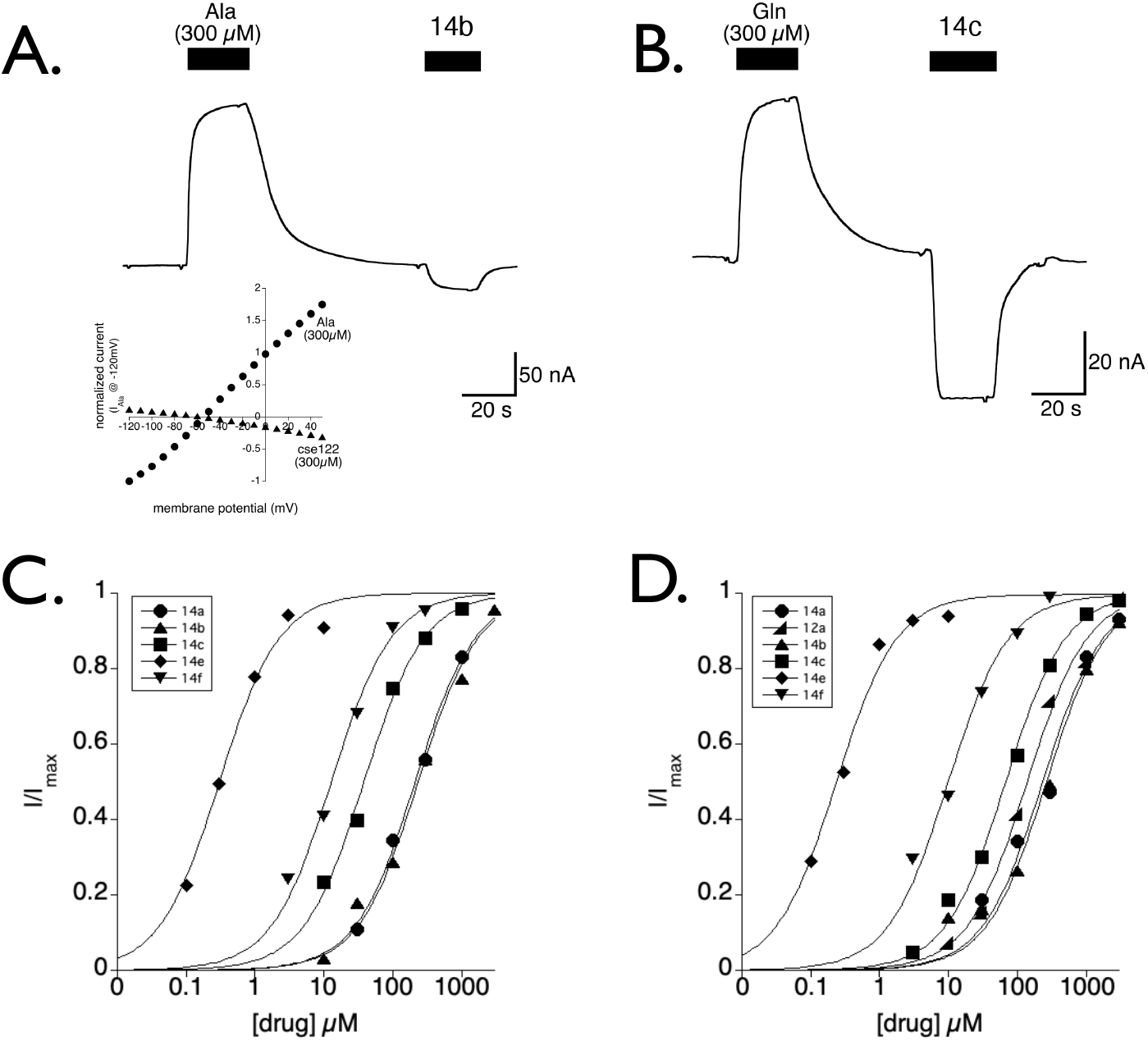
Hydroxy proline derivatives block a leak current in SLC1A4 and mSLC1A5. (**A &B)**. Representative traces showing the anion current gated by substrates (alanine and glutamine) or blocked by the prolinol derivative 14b and 14c for SLC1A4 and m SLC1A5, respectively. *Inset A:* The current and voltage relationship for SLC1A4 currents activated by Ala or blocked by 14b resolved by subtracting away voltage jumps in buffer. (**C&D)**. The leak current in SLC1A4 and mSLC1A5 has a saturable block revealed by increasing concentrations of 14e (down triangle), 14f (diamond), 14b (square), 14c (up triangle), 12a (wedge), and 14a (circles). The 50% block is considered to be the Ki for binding of the non-transportable substrate.

**Table 1.**
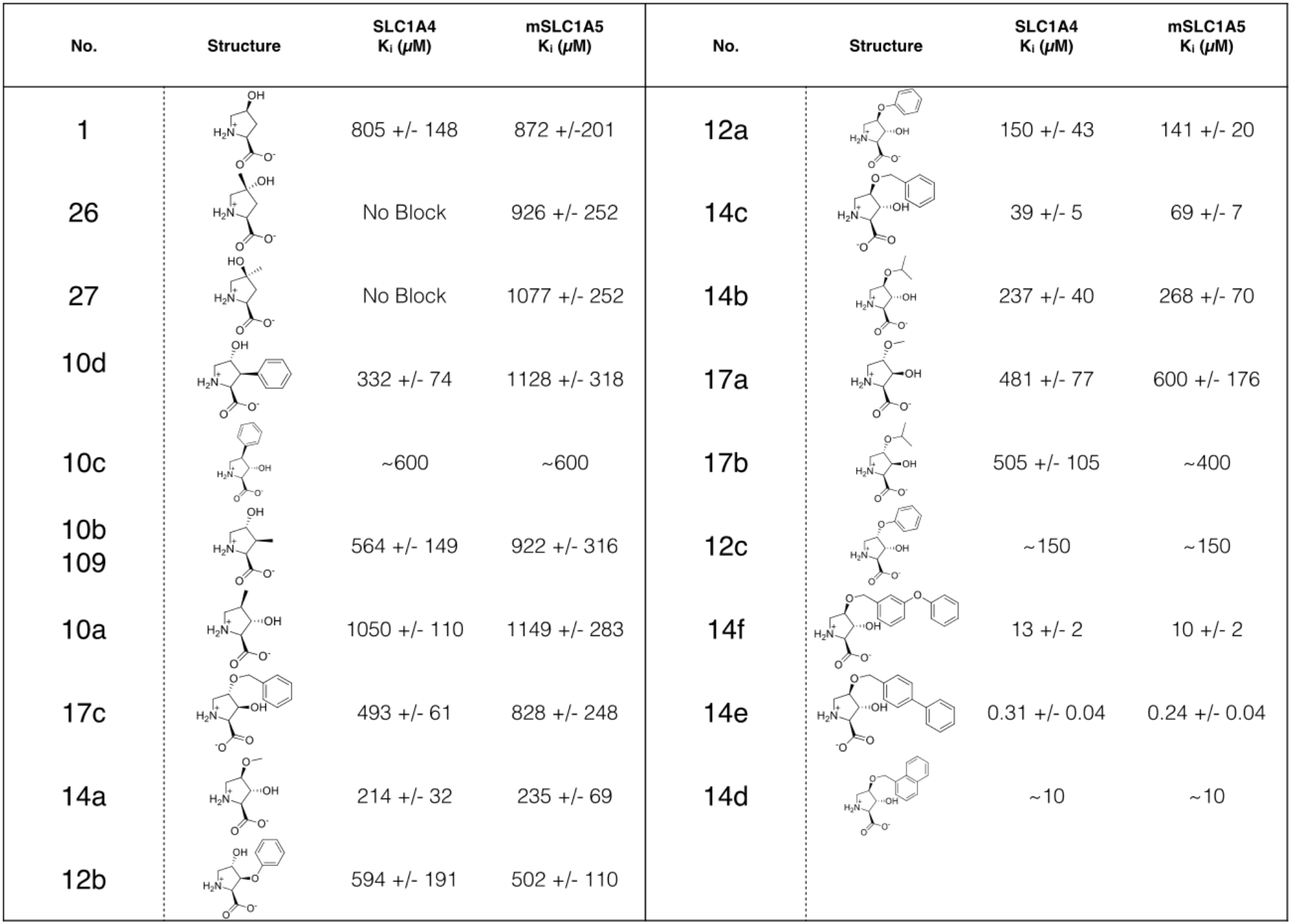
Prolinol K_i_ values for SLC1A4 and mSLClA5. Values were calculated from the dose response of blocked anion conductance. Compounds tested only once are shown without standard error.

#### BPOHP Pharmacology

We further characterized the lead compound from this study, biphenyl-O-hydroxy-proline (BPOHP, **14e**). Radiolabeled [^3^H]L-alanine (1uM) was used to monitor both uptake as well as transporter exchange in SLC1A4. BPOHP blocked [^3^H]L-alanine uptake with a slightly lower yet comparable affinity determined from inhibition of the anion leak current (Figure 4A: IC50 = 128nM). As obligate exchangers, SLC1A4 transporters require intracellular substrate binding and translocation to reset the transporter extracellular binding site. In an effort to determine whether BPOHP was a substrate, we preloaded [^3^H]L-alanine into oocytes expressing SLC1A4 or mSLC1A5, and then monitored the ability of buffer, 1mM L-alanine, or 3uM BPOHP to induce exchange of intracellular [^3^H]L-alanine. The results demonstrated a clear ability for L-alanine but not buffer nor BPOHP to induce exchange, supporting this series of compounds as non-transported inhibitors (Figure 4B). To investigate the nature of BPOHP competitive binding at SLC1A4, we measured L-alanine dose responses in the presence and absence of BPOHP. A respective right shift in the substrate dose response was apparent with increasing concentrations of BPOHP (0.1, 0.3, and 1uM_Figure 4C). A Schild analysis, plotting the log plot of the dose ratio vs a log of inhibitor concentration, revealed a clear linear relationship, characteristic of competitive inhibition (Figure 4D, K_B_=120nM).

**Figure 4.**
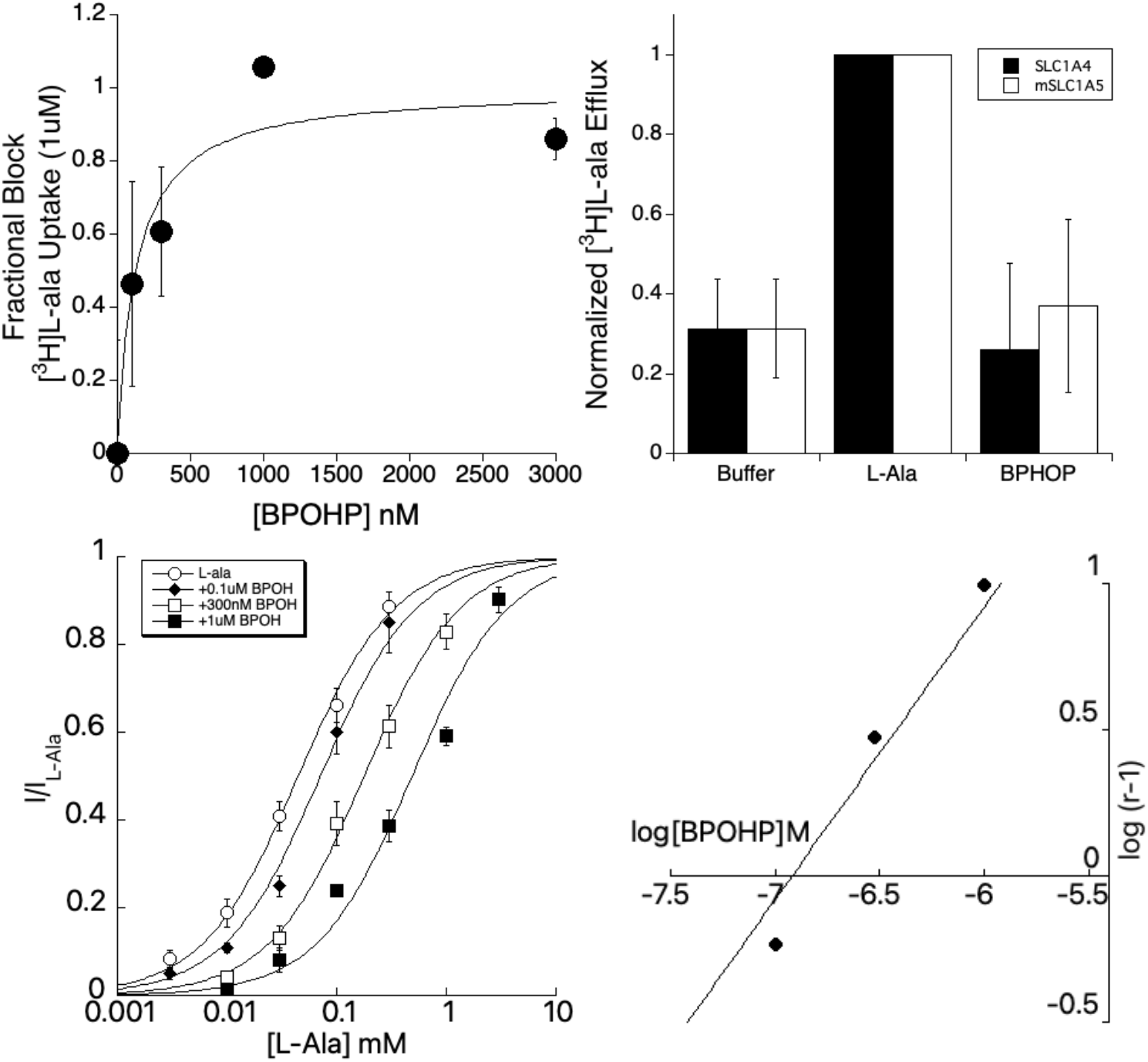
BPOHP (14e) characterization as a competitive, non transported inhibitor. **A**. SLC1A4 injected oocytes have tritiated [^3^H]L-ala uptake blocked dose dependently by BPOHP (IC = 128 nM). **B**. Radiolabeled [^3^H]L-ala exchange for SLC1A4 and m SLC1A5. Exchange was normalized to the amount induced by 1mM L-ala. No exchange was apparent for 3uM BPOHP. **C**. L-ala dose response curves for SLC1A4, showing the right shift with increasing concentrations of BPHOP (0, 0.1, 0.3, 1 uM). **D**. A Schild plot showing a linear relationship between the dose ratio and the concentrations of BPOHP, confirming competitive inhibition.

Cross reactivity of BPOHP experiments were performed against glutamate transporters (SLC1A 1-3) within the SLC1A gene family as well as at neutral amino acid transporters that have overlapping substrate profiles, SLC38A1,2, and 4 transporters (ATA1, ATA2, ATA3) and SLC7A10+SLC3A2 (asc-1/4f2). Using TEVC, BPOHP did block a 10uM glutamate current at higher concentrations with the largest overlap with EAAT2 (Supplemental Fig. 2_IC50 for EAAT1 = 2.45uM, EAAT2, 1.2uM, and EAAT3 = 25.3 uM). Oocytes expressing SLC38A1,2, and 4 transporters produce electrogenic coupled currents in response to L-alanine (100uM); these currents showed no discernable block at 3uM BPOHP. Finally, radiolabeled [^3^H]D-serine uptake was used to assess the electroneutral SLC7A10+SLC3A2 transporter, in which 10uM BPOHP had no effect (Supplemental Figure 3).

### Molecular Docking

Computational docking predicts prolinol orientation within the substrate ligand binding pocket. We analyzed our synthetic prolinol series via computational docking using a homologymodel of human SLC1A5 (ASCT2) derived from a ligand bound (TBOA) co-crystal structure of Glt_Ph_, 2NWW.^23^ Our initial computational docking was performed with one parameter constraint in favor of a polar interaction with aspartate residue D476 of SLC1A5. With this constraint, Golds scoring function accurately predicted the relative SLC1A5 inhibitor activity in rank order for the pyrrolidine derivatives as we have found empirically.

Prolinol **14c** (purple) docked into the SLC1A5 homology model by aligning its secondary amine to the *β*-carboxylic acid of D476 on transmembrane 8 (TM8, green) at 2.2Å and to the nearby T480 (Figure 5A). The *α*-carboxylic acid of **14c** is oriented towards the amide of nearby N483 on TM8 and serine peptide amide of the HP1 loop (blue). Moreover, the (3*R*) alcohol of the prolinol **14c** aligns near the sulfhydryl of C479. The closed ring limits the confomraitnal space assumed by the pyrrolidine ring, and this orientation places the benzylether moiety of **14c** between the membrane spanning side of helix transmembrane seven (TM7a; teal) and HP2 loop (red). The benzyloxy is positioned over methionine residue, M399 on TM7, which mimics the benzyloxy substituent of TBOA over methionine M311 of Glt_Ph_, 2NWW.^23^ In contrast, the functionally inactive isomer **17c**, *cis*-3-hydroxy-*trans*-4-benzyloxy-L-proline (yellow), does not allow for a polar interaction with C479 even though the benzylic ether orientates within a plausible hydrophobic pocket perpendicular the phenyl ring of F405 (Figure 5A).^30^ Docking of **12c** into the homology model (not represented) positions the phenoxy substituent adjacent to the phenyl ring of F405 like **17c** but positioning (3*R*) alcohol, like **14c**, to interact with C479. The empirical results validate the benefit of both the C479 contact with the trans-3-hydroxyl as well as a hydrophobic binding pocket near F405.

**Figure 5.**
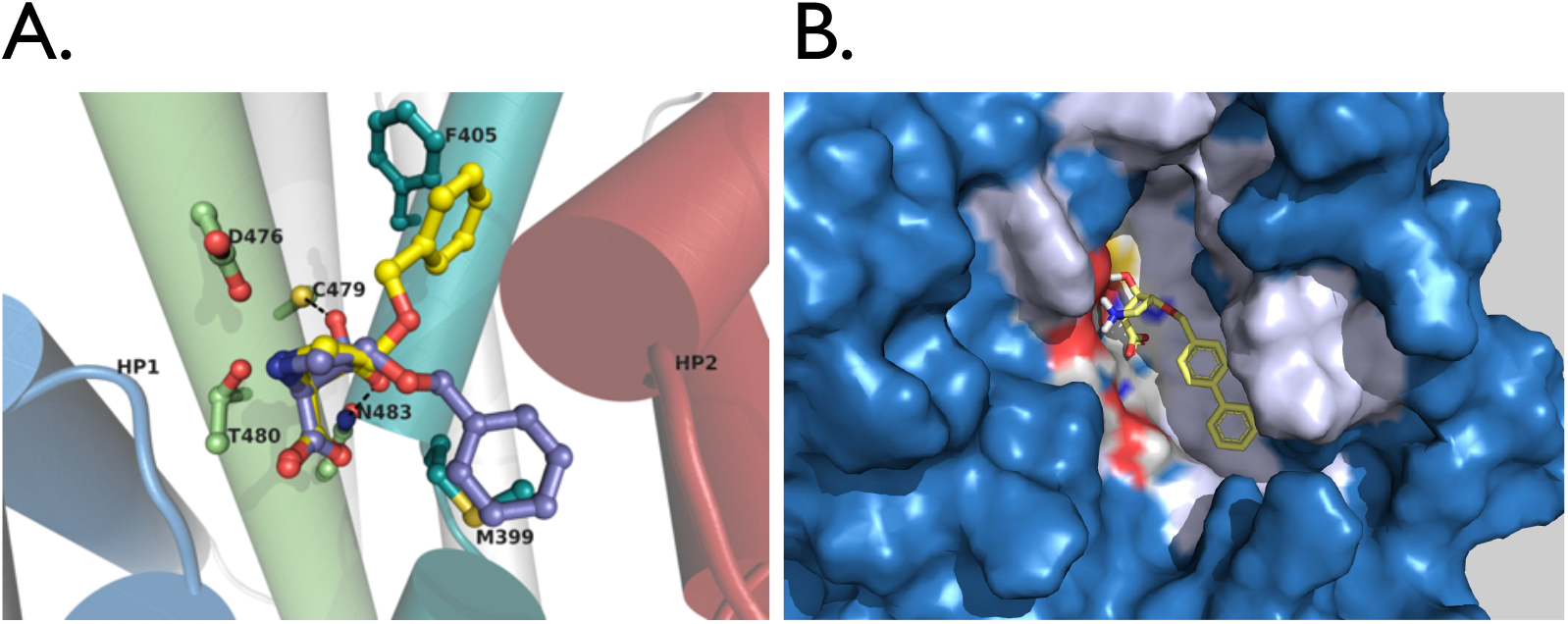
Molecular Docking. **A**. Prolinol **17c** (yellow) and **14c** (purple) overlain in an ASCT2 homology model illustrating two probable hydrophobic pockets, one near phenylalanine F405 and the other on top of a methionine residue (M399) on TMD7 (teal). **B**. A surface rendering of the lead compound **14e** docked into the SLC1A5 homology model binding site.

Docking of our most potent lead **14e** predicts a hydrophobic pocket that extends from M399 and along TM7α (Fig. 5B). The orientation and position of **14e** is consistent with that predicted in our docking solutions with **14c** where again the (3*R*) alcohol is in contact with C479, the pyrrolidine amine is positioned in contact between D476 and T480, and finally the *α*-carboxylic acid interacting with N483 and serine *α*-carbon peptide backbone of HP1. The larger arene substituted prolinols,**14d-f**, score relatively better than the benzylic-ether substituent of 14c, which is also refected in the increase in affinity.

## Discussion

*Trans*-4-hydroxy-proline (4-HP), a previously identified substrate for SLC1A5 (ASCT2),^22^ led to our identification of *trans*-3-hydroxy-proline (3-HP) as a substrate of SLC1A4 and SLC1A5. Using these two conformationally constrained scaffolds, a unique library of compounds evolved to elucidate critical molecular orientations that select for SLC1A4 and SLC1A5 binding. Through electrophysiology, radiolabel transport assays, and molecular docking, this series of inhibitors was fine-tuned through three iterations of chemical design and expansion. Cross reactivity was assessed on glutamate transporters of the SLC1A gene family in addition to neutral amino acid transporters with similar amino acid profiles and tissue distribution (SLC38A1 (ATA1), SLC38A2 (ATA2), SLC38A3 (ATA3) and SLC7A10 (asc-1) +SLC3A2 (4f2). The lead compound, BPOHP, marks an advancement in SLC1A4/5 pharmacology with a Ki = 120nM and will aid in directing future ligand design to obtain high resolution co-crystal structures.

Due to the convenience of commercial availability, the entire prolinol series developed within this study was derived from *trans*-4-hydroxy-L-proline. Our first active ‘hit’ in this series towards SLC1A4/5, though with modest affinity, was *trans*-3-hydroxy-*cis*-4-methoxy-L-proline **14a**. Surprisingly, the methyl analogue, *trans*-3-hydroxy-*cis*-4-methyl-L-proline **10a**, showed no inhibition of transport. A possible explanation for the observed loss in activity may be that these two prolinols assume significantly rigid and dissimilar boat conformations, supported by energy minimization and docking. Extension of **14a** to produce the isopropoxy-prolinol **14b** provided no additional benefit in activity. Introduction of a phenoxy to produce **12a** did add a measurable increase in affinity. However, the greatest increase in potency came by extension of the methoxy substituent to arrive at **14c**, a benzyloxy derivative. From our docking, it is not exactly clear as to why the benzyloxy substitution affords greater activity than the phenol ether **12a**, though, it may stem from the added conformational flexibility of the benzyloxy group. This may allow the benzyl group to position slightly underneath the HP2 loop adjacent to the hydrophobic residue M399, like the orientation of TBOA in the Glt_Ph_ structure.^23^

Considering the optimization of EAAT inhibitors, the extension of the benzyloxy moiety of TBOA led to the discovery of TFB-TBOA, an inhibitor with over a 250-fold improved potency.^31^ Similarly the numerous analogs of aryl-aspartamides in which substitution of a phenyl amide group for a fluorene or a biphenyl substituent noticeably increased potency at glutamate transporters.^30,32^ Considering SLC1A4/5 an increase in potency also is observed by extension of benzyl-ether derivatives of **14c** including biphenyl, naphthyl or diphenylether containing analogs. Of the three tested biphenyl ether (2S, 3R, 4R)-3-hydroxy-4-O-methyl-p-biphenyl-proline, BPOHP (**14e)** was the most potent inhibitor at both SLC1A4/5 transporters. The extended hydrophobic pocket between HP2 and TM7 may accommodate benzylic ether analogues of 14d-f of greater steric size and diversity, e.g., halogen, methyl, methoxy substituted biphenyls, the structurally rigid aromatic fluorene or substituting with heteroarenes. Our docking results preicted that the hydrophobic pocket near F405 confers high affinity binding within our SLC1A5 homology model. Considering that the hydrophobic pocket adjacent to HP2 over F405 is only modestly conserved among the various transporter subtypes of the SLC1 family reveals several implications for future ligand design stratagies.

Although incomplete, this prolinol series has begun to describe a structure-activity relationship (SAR) for designing non-substrate inhibitors of the SLC1A4/5 transporters and may also provide an entry to design a SLC1A4/5 subtype selective inhibitor. Glutamine gives one point of separation in drug design between SLC1A4 and SLC1A5. Structurally the R447 residue that confers selectivity for acid amino acid substrates of glutamate transporters also distinguishes glutamine selectivity between the SLC1A4 and SLC1A5.^33,34^ It is worth noting a second mutation in TMD8 was needed to confer Gln affinity in SLC1A4 which the authors hypothesize is a size constraint of the binding pockets.^34^ One interesting avenue to pursue these unique molecular contacts for SLC1A4 inhibitors could include analogs of *cis*-3-phenyl-4-*trans*-hydroxy-L-proline, **10d**. The 3-positioned *cis*-methyl prolinol **10b** from our screening demonstrated slight selectivity for SLC1A4 over SLC1A5. The selectivity is further improved with a phenyl substituent, **10d**, suggested by greater potency at SLC1A4 and no detectable change at SLC1A5. Furthermore, these observations will inform our ligand design strategies to discover subtype selective inhibitors for the SLC1A transporters.

High resolution structural models for SLC1A4 and SLC1A5 have been developed with cryo-EM.^35–37^ These structures confirm SLC1A4/5 homology modeling of acharel/glutatmate transporters offering insight into variations that distinguish obligate exchange, couple transport, ion coordination, inhibitor binding, and monogenic disorders. Within the SLC1A gene family, the most potent competitive inhibitors assume a similar orientation to TBOA in the original Glt_Ph_ crystal structures as does BPOHP docked into Glt_Ph_ (Figure 5B).^23,38^ Alternative binding pockets for SLC1A5 inhibitors have been described to occupy the hydrophobic groove between TMD7 and HP2^34,39^, however, like **17c and 12c**, which may orientate into this TMD7/HP2 pocket, these inhibitors exhibit a substantially lower affinity for SLC1A4/5. While writing this manuscript Garibsingh et al. 2021 published work on a proline scaffolds that generated molecules of similar design in combination with some low resolution structures that helps validate our homology model and chemistry evolution.^40^ We will emphasize that our work on prolinols predates their molecular docking publications of prolinols, but has not yet been recognized in the historical scientific literature though clearly documented in the public domain (Patent US 20130065935A1). Of importance, both works demonstrate an improvement in drug design for SLC1A4/5 that has had no molecules with affinity in the nanomlar range. Specifically, our lead compounds were built off the *trans*-3-hydroxy scaffold and reach a 7-25 fold improvement in affinity.^20,40^ The (3*R*) alcohol provides a beneficial contact with C479 allowing for the increased potency observed for BPOHP. BPOHP dose exhibit some cross reactivity at glutamate transporters with an approximately 8-10 fold lower affinity. No overlap in inhibition was measured at neutral amino acid transporters from the SLC38 gene family (SLC38A1, SLC38A2, SLC38A4) or at SLC7A10 (asc-1) +SLC3A2 (4f2) transporter (Supplemental Fig. 3).

Altering glutamine equilibrium in tumor cells has been an effective strategy in limiting proliferation, and SLC1A5 represents a potential target to achieve this aim.^41^ Multiple constrained analogs based on substrates have been developed with the intention of selectively inhibiting SLC1A5 as a tumor-suppressing strategy, but documented cross-reactivity still obscures our understanding of the specific role SLC1A5 in tumor growth.^42–45^ These examples emphasize the importance of controlling and defining cross-reactivity in the pharmacological characterization of lead compounds. Considering ligands based on the prolinol scaffold, future efforts may focus on optimizing selectivity over the other transporters of the SLC1A gene family because of structurally conserved binding pockets. Assuming poor blood-barrier permeability, the overlap with glutamate transporters described in this work may not be as detrimental in peripheral tissues where glutamate transporters are expressed at a much lower density than within the CNS.^46^ Other transporters that should be considered for inhibitor cross reactivity include the neutral amino acid transporters from the SLC38A and SLC7A+SLC3A2 gene families.

## Supporting information

Supplemental Results

## Ancillary Information

*Supplemental Information and Expermientals are reported in a separate document*.

## Acknowledgment

We recognize and remember C. Sean Esslinger for his work on two fundamental lead compounds developed for altering glutamine and SLC1A5. His mentoring and cleverness in chemical design led to both the GNPA compounds and these stereoselective prolinol derivatives.

This work was supported by the National Institutes of Health grant R01 MH110646.

## Notes

### Competing Interest Statement

The authors have declared no competing interest.

### Summary of Updates

Grammatical edits.

