## Supplemental Results for "Discovery and synthesis of hydroxy-L-proline based blockers of the neutral amino acid transporters SLC1A4 (ASCT1) and SLC1A5 (ASCT2)"

### Supplementary Figures

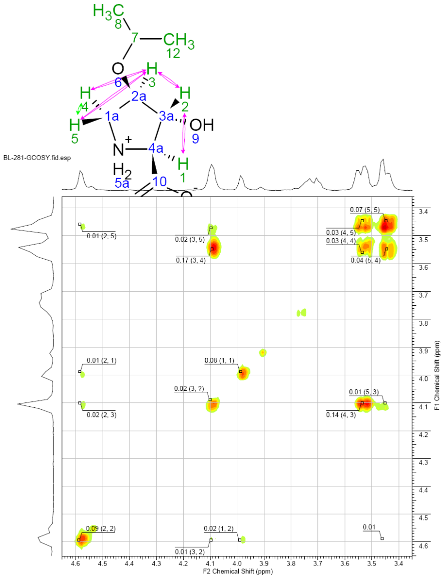


**S.Figure 1** gCOSY of 14b (CSE122), shows J-coupling correlation as intensity in 0.0x and proton # in (x,x). The numerical labels such as 0.09 (2,2) indicate intensity (first number) and protons correlated in parentheses. The methylene protons H-4 & H-5 are easily spotted at ppm 3.4-3.56 by their strong J2 coupling and chemical shift. It is clear that the methylene protons are coupled to a proton at ppm 4.1. Importantly, one of these methylene protons, at 3.56 ppm, is strongly coupled to this proton, possibly a result of a small dihedral angle (0-20°) between two *cis* vicinal protons suggesting they reside on the same orientation of the pyrrolidine ring. A 2D NOESY experiment confirms these vicinal proton nuclei reside at a respective *cis* orientation by exhibiting an NOE enhancement. Additionally, an NOE enhancement is not observed between the other methylene proton at 3.42 ppm and the proton at 4.1 ppm. The alpha proton H-1, seen at ppm 3.98, is weakly coupled to only one other proton (assigned H-2) at ppm 4.58, and exhibited no detectable NOE with this proton suggesting they reside on opposite planes or respective *trans* orientation. This proton (H-2) has small but detectable couplings with two other protons at 3.42 (a methylene proton) and at 4.1 ppm. The weak intensity of the J3 couplings and its chemical shift suggest this proton is H-2 and therefore the proton at ppm 4.1 is H-3. The methylene protons were then assigned according to coupling with the H-3 proton, with H-4 at ppm 3.56 and H-5 at ppm 3.42. This assignment is also consistent with the small J4 coupling observed between H-2 and H-5.

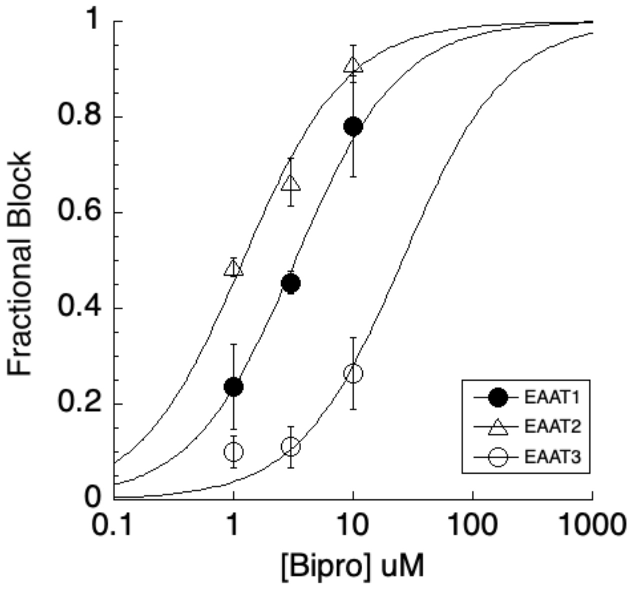


**S.Figure 2 A.** TEVC from oocytes expressing SLC1A1-3 transporters. Plot illustrates the fractional block of the 10uM L-glu current by increasing concentrations of BPOHP. The IC50 values are SLC1A3_filled circle (EAAT1) = 1.75uM, SLC1A2_triangle (EAAT2) = 1.2 uM, and SLC1A1_open circle (EAAT3) = 19.5 uM.

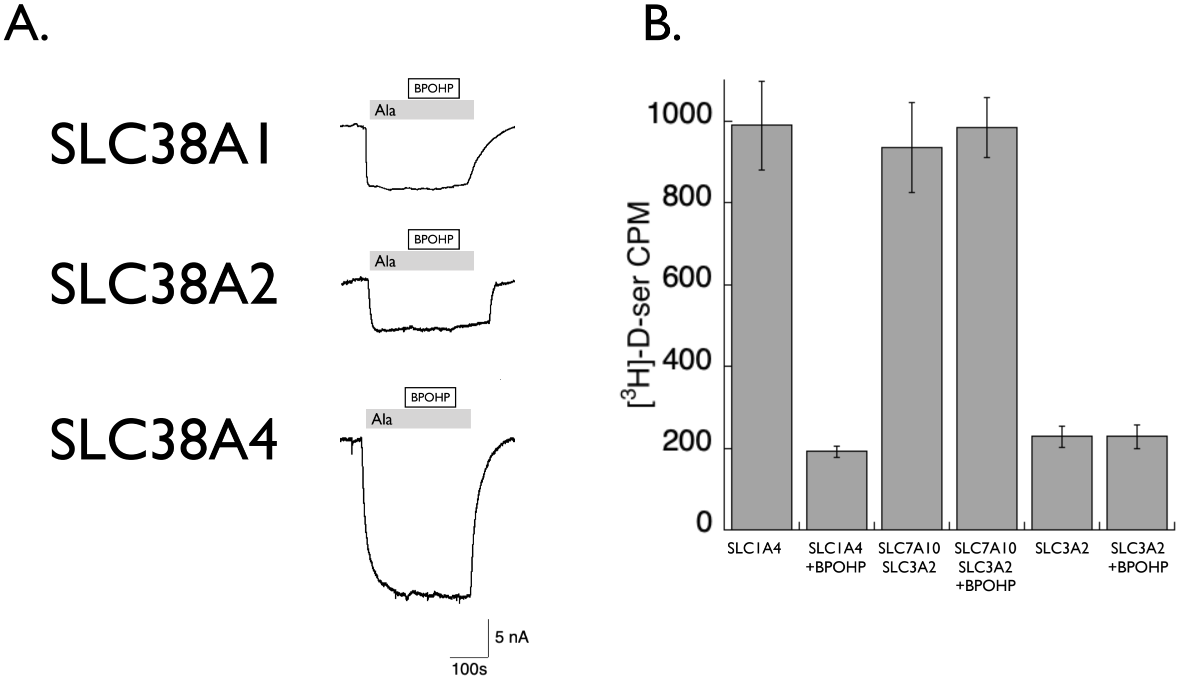


S.**Figure 3** **A.** Representative traces showing 100uM L-Ala currents and co-application of 3uM BPOHP (black bar) with no effect on SLC38A neutral amino acid transporters SLC38A1(ATA1), SLC38A2 (ATA2), and SLC38A4 (ATA3). **B.** [$^3$H]D-serine uptake (100nM) in SLC1A4 is blocked by BPOHP (10uM), but not the $H^3$D-serine uptake by SLC7A10+SLC3A2 (asc1+4F2). Also no effect was seen for SLC3A2 (4f2) injected alone.

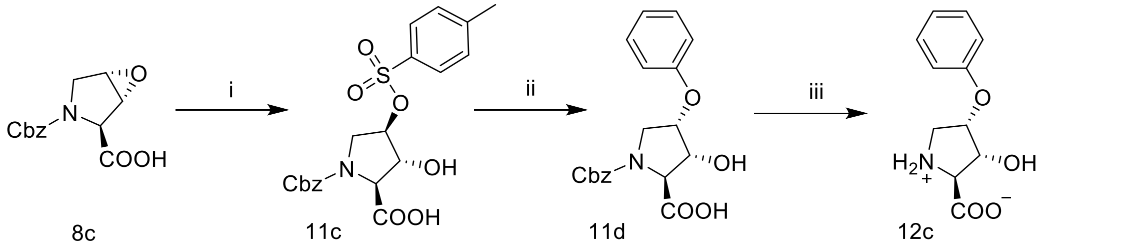


**S. Figure 4**. (i) Tosylic acid 0.1M in THF 60°C. (ii) Phenol and NaH in DCM, 40°C under argon balloon (iii) 10% Pd/C (10% w/eq.), $H_2$, MeOH, $H_2O$, $^+NH_4^-OAc$ (0.5 eq).

#### Experimental Procedures

Proton $^1H$ and carbon $^{13}C$ nuclear magnetic resonance (NMR) spectra were obtained from Varian 500 MHz spectrometers as specified. Both $^1H$ and $^{13}C$ spectra are reported in parts per million (ppm or $\delta$) downfield from tetramethylsilane as an internal standard. All chemical shifts reported were referenced to residual protonated $D_2O$ or DMSO solvent peaks at 4.75 or 2.5 ppm, respectively, for $^1H$ proton spectra and 39.5 ppm for $^{13}C$ carbon spectra unless specified otherwise. Multiplicity is reported as doublets (d), doublet of doublets (dd) etc. where $\Sigma$ represents the sum of all couplings, b = broad. N.O.e. experiments were performed on a Varian 500 MHz using 2D NOESY parameters with zero-quantum suppression and mixing times of ~0.7 s to aid in making stereo and regiochemistry assignments. Gradient correlation spectroscopy (gCOSY) to aid in determining regiochemistry was also performed using a Varian 500 MHz. Optical rotations of final compounds dissolved in methanol or DMSO, as specified, with concentrations given in g/100 mL were determined via a Perkin-Elmer Model 241 polarimeter in a 1.0 dm glass cell sodium D line. High resolution mass spectra (HRMS) were determined on a Waters LC-MS spectrometer using caffeine or trimethoprim as a mass standard at m/z 195.0882 or 291.1457 respectively for HRMS. All reagents were purchased from Acros Organics and used without further purification. Solvents as well as 40-60 μM silica gel for flash chromatography were obtained from EMD chemicals (Gibbstown, NJ). Intermediates were isolated by silica gel used without further purification, unless specified otherwise. A mobile phase consisting of 95% $CH_2Cl_2$ / 4% MeOH / 1% AcOH or hexanes / ethyl acetate was used for TLC and flash chromatography, as specified. Ion exchange chromatography was carried out using Bio-rad AG® 50W cation exchange resin which was pre-equilibrated in 1M HCl followed by deionized water. Semi-prep reverse phase

HPLC was carried out using a C18, 250 mm x 21.2 mm, 10 μ, Chromegabond WR 120A column from ES Industries using acetonitrile / water 0.5% ammonium acetate.

*N-Cbz-4-hydroxyproline* **3**. *trans*-4-Hydroxyproline **2** (27.5 g, 209.7 mmol) was dissolved into 200mL of 50% MeOH in water. While on ice bath, to the mixture was added 85.9 mL of benzyl chloroformate (0.252 mmol; 1.2 eq; 50% vol. in toluene) dropwise. Pellets of NaOH were added to maintain the pH at ~9-10. The mixture was then allowed to warm to rt (22˚C). The mixture was allowed to stir over night at rt and the pH closely monitored and maintained at pH = 10 by addition of NaOH pellets. The mixture was then diluted by addition of 200 mL water and then extracted with 3 x 200 mL toluene (to remove remaining benzyl chloroformate). The aqueous layer was then salted with NaCl until saturated and the pH was adjusted to ~2 by addition of HCl conc. This was then extracted with 4 x 200 mL of ethyl acetate. The ethyl acetate washes were then combined and concentrated via rotovap, leaving an oil residue. Dichloromethane (50 mL) was added 4 times to remaining oil and then removed by vacuum. Hexanes (50 mL) was then added to the oil and again removed by vacuum, leaving a sticky foam N-Cbz-4-hydroxyproline crude (quantitative yield) which was then used w/o any further purification. This residue can also be recrystallized in 10:1, hexanes:$CH_2Cl_2$. $^1H$ and $^{13}C$ NMR spectra were in agreement with $^1H$ and $^{13}C$ NMR spectra previously reported [Robinson; *et al.* 1998].

*N-Cbz-4-hydroxyproline benzyl ester* **4**. N-Cbz-4-hydroxyproline **3** (40.00 g, 150.8 mmol) was dissolved into 110 mL DMF. To this mixture was added 45.85 g of $K_2CO_3$ (2.2 eq; 331.7 mmol) and 2.30 g of NaI (0.1 eq). The reaction flask was purged with argon and 55.4 mL of benzylbromide (3 eq; 452 mmol) was added dropwise. This mixture was then stirred over night at rt. The reaction was then diluted with 300 mL ethyl acetate and washed with water (8 x 200 mL) to remove DMF, followed by washing with 100 mL brine. The organic layer was dried over $Na_2SO_4$ and concentrated to a yellow oil. The oil was then washed several times with 100 mL hexanes to remove any remaining benzyl bromide and then separated by silica gel in hexanes : ethyl acetate, 3:2, isolating N-Cbz-4-hydroxyproline benzyl ester **4** at $R_f$ = 0.25 in nearly quantitative yield. $^{1}H$ and $^{13}C$ NMR spectra were in agreement with $^{1}H$ and $^{13}C$ NMR spectra previously reported [Robinson; *et al.* 1998].

*N-Cbz-4-p-toluenesulfonyl-ether-proline-benzyl ester* **5**. N-Cbz-4-hydroxyproline benzyl ester **4** (21.77 g, 61.3 mmol) was dissolved into 150 mL cold pyridine 0˚C. Immediately, 12.976 g of p-toluenesulfonyl chloride (1.1 eq.; 67.4 mmol) was added. The flask containing this mixture was then purged with argon and placed into a refrigerator at ~0˚C for one full week with occasional vortex. After one week it was observed that a significant amount of pyridine-HCl had formed and all the starting material appeared to have been consumed (determined by TLC). The mixture was then taken up into 100 mL $Et_2O$ and 50 mL ethyl acetate and washed 2 x 100 mL cold 5% HCl and then 3 x 200 mL water. The organic layers were combined and dried over $Na_2SO_4$, filtered and concentrated to an oil. The product N-Cbz-4-p-toluenesulfonyl-ether-proline-benzyl ester **5** was isolated by silica gel in hexanes / ethyl acetate, 7:3; $R_f$ = 0.25 to recover 28.05 g, (90% yield). $^{1}H$ and $^{13}C$ NMR spectra were in agreement with $^{1}H$ and $^{13}C$ NMR spectra previously reported [Robinson; *et al*. 1998].

*N-Cbz-4-phenylseleno-L-proline-benzyl ester* **6**. Diphenyldiselenide (9.504 g, 0.6 eq.; 30.2 mmol) was dissolved into 60 mL of tBuOH and purged of air by vacuum and replaced with argon. The solution was brought to a reflux where of $NaBH_4$ (2.293 g, 1.2 eq, 60.3 mmol) was added slowly over 15 min. This mixture was refluxed for 30 min until the mixture had turned completely white. To this was added N-Cbz-4-p-toluenesulfonyl-ether-proline-benzyl ester **5** (25.6 g, 50.2 mmol) in 40mL tBuOH. The reflux was continued under argon for ~3 h then allowed to cool to rt. The mixture was then diluted with 200 mL ethyl acetate and washed 3 x 250 mL water followed by 100 mL brine then treated with $MgSO_4$ to dry, filtered and concentrated to a crude oil. The product N-Cbz-4-phenylseleno-ʟ-proline-benzyl ester **6** was isolated by silica gel 3:1 hexanes : ethyl acetate at $R_f$ = 0.3, recovering 22.21 g, (89% yield). $^1H$ and $^{13}C$ NMR spectra were in agreement with $^1H$ and $^{13}C$ NMR spectra previously reported [Robinson; *et al.* 1998].

*N-Cbz-3,4-dehydro-ʟ-proline-benzyl ester* **7**. N-Cbz-4-phenylseleno-ʟ-proline-benzyl ester **6** (22.11 g, 44.7 mmol) was dissolved in 200 mL $CH_2Cl_2$ (warning! explosive conditions if concentrated: needs to be performed in more dilute conditions than in this example or ~200 mL per 10 g of starting material). The solution was chilled on ice bath over argon and 4.9 mL pyridine (1.34 eq.; 60 mmol) was added followed by addition of 11.50 mL 30% $H_2O_2$ (2.5 eq.; 112.0 mmol) drop wise. The solution was gradually allowed to return to room temp over the course of one hour where it was stirred for 2.5 h. The solution, now brown in color, was diluted with 100 mL $CH_2Cl_2$ and washed with 100 mL of 5% $NaHSO_4$, 2 x 50 mL $NaHCO_3$, and 3 x 150 mL water. The organic layer was dried over $Na_2SO_4$ and filtered. A total of 12.02 g of N-Cbz-3,4-dehydro-ʟ-proline-benzyl ester **7** was isolated by silica gel in mobile phase of 4:1 hexanes / etheyl acetate; $R_f$ = 0.15; (80% yield). $^1H$ and $^{13}C$ NMR spectra were in agreement with $^1H$ and $^{13}C$ NMR spectra previously reported [Robinson; *et al.* 1998].

*N-Cbz-trans-3,4-epoxy-L-proline-benzyl ester **8a** and N-Cbz-cis-3,4-epoxy-L-proline-benzyl ester **8b***. N-Cbz-3,4-dehydro-L-proline-benzyl ester (9.80 g, 29.0 mmol) **7** was dissolved into 12 mL $CH_2Cl_2$. To this was added mCPBA (14.3 g, 70% weight; 58.1 mmol; 2 eq.) and 2,8-di-tert-butyl-4-methylphenol (30 mg, radical inhibitor). The solution was heated to reflux and allowed to stir overnight under argon balloon. An additional 1 eq. of mCPBA (7.15 g) was added and the solution was allowed to reflux again overnight. The flask was then placed on an ice bath where a white precipitate formed and was subsequently filtered using cold dichloromethane to wash. The $CH_2Cl_2$ was removed by rotovap leaving a white solid residue. The residue was taken up into 150 mL $Et_2O$ and washed 8 times with 100 mL (aq) $NaHCO_3$, 2 x 100 mL water, and 100 mL brine then dried over $Na_2SO_4$ and filtered. The remaining oil was then separated by silica gel in 4:1 hexanes / ethyl acetate. The *trans* isomer 5.228 g, N-Cbz-3,4-*trans*-epoxy-L-proline-benzyl ester **8a**, was isolated at $R_f$ = 0.2 and the *cis* isomer 4.100 g, N-Cbz-3,4-*cis*-epoxy-L-proline-benzyl ester **8b** being isolated at $R_f$ = 0.1, a total yield of 91% and ratio of 5:4 trans / cis. $^1$H and $^{13}$C NMR spectra for both diastereomers were in agreement with $^1$H and $^{13}$C NMR spectra previously reported [Robinson; *et al.* 1998].

*N-Cbz-trans-4-methylxanthate-L-proline* **4a**. To 3.635 g (13.7 mmol) of N-Cbz-*trans*-4-hydroxy-L-proline **3** was added 35 mL of dry THF and slow addition of 1.73 g of NaH (95%; 5 eq.; 68.5 mmol). The solution was then purged with argon and allowed to stir for 10 min. To this was added 4.14 mL of $CS_2$ (68.5 mmol; 5 eq.) and allowed to stir for 2 h at rt, forming the xanthate salt. To the flask was then added 2.87 mL of iodomethane (48.0 mmol; 3.5 eq) and allowed to stir overnight at rt and under argon balloon. The reaction mixture was then chilled on ice bath where ~10 mL of glacial acetic acid was added to quench the excess NaH (heavy white precipitate forms). The quenched reaction mixture was then diluted with 100 mL of $Et_2O$, filtered using excess ether, and then concentrated under reduced pressure, leaving a yellow / brown oil. The final product, 4.19 g, N-Cbz-*trans*-4-methylxanthate-ʟ-proline **4a** was isolated by silica gel at $R_f$ = 0.15 in 97% $CH_2Cl_2$, 2% MeOH 1% AcOH, 70% yield as a crude mixture which was used in the next step without further purification.

*N-Cbz-trans-4-methylxanthate-ʟ-proline benzyl ester* **5a**. To 3.40 g (9.57 mmol) of N-Cbz-*trans*-4-methylxanthate-ʟ-proline **4a** was added 30 mL DMF, 2.64 g (19.1 mmol; 2 eq.) of $K_2CO_3$, and 143 mg of NaI (0.957 mmol; 0.1 eq.). The reaction flask was purged with argon, and 3.58 mL of benzyl bromide (98 %; 29.3 mmol; 3 eq). The mixture was allowed to stir at rt for 24 h under argon balloon. The solution was taken up into 100 mL ethyl acetate and then washed with $H_2O$ (4 x 300 mL). The organic layer was separated and dried with $MgSO_4$, filtered and concentrated to a yellow oil. The final product N-Cbz-*trans*-4-methylxanthate-ʟ-proline benzyl ester **5a** was isolated by silica gel in 4: 1 hexanes / ethyl acetate, $R_f$ = 0.25, with a quantitative yield. $^1$H-NMR (400 MHz, $CDCl_3$) δ 7.40-7.20 (m, 10H), 5.95 (d, J = 2.2 Hz, 1H), 5.25-5.16 (m, 1H), 5.17 (s, 1H), 5.07 (s, 1H), 5.00 (s, 1H), 4.62-4.52 (ddd, J = $44.0_{\Sigma}$, 28.6, 8.1 Hz, 1H), 3.99 (d, J = 12.4 Hz, 1H), 3.88 (s, 1H), 2.67-2.59 (m, J = 8.1, 2.2 Hz, 1H), 2.53 (s, 3H), 2.36 (m, J = 8.1, 2.1 Hz, 1H).

*N-Cbz-3,4-dehydro-ʟ-proline* **7a**. An open flask containing 0.240 g of N-Cbz-*trans*-4-methylxanthate-ʟ-proline (0.675 mmol) **4a**, 1 mL ethanol, 10 mL water, and 42 mg $Na_2CO_3$ (anhydrous; 0.5 eq) was placed into a microwave on medium (100 watt noncontinous radiation; retail purchased Sunbeam microwave) for 4 min. The contents were allowed to cool to rt (keep in hood; xanthate elimination liberates $CS_2$ and other methyl sulfides, producing a very undesirable noxious smell). The reaction mixture was diluted with 20 mL water and the pH was adjusted to ~2-3 by addition of $H_3PO_4$. The aqueous mixture was salted with NaCl until saturated, then extracted with 3 x 50 mL ethyl acetate, dried with $Na_2SO_4$, and concentrated to a crude orange colored oil. The product N-Cbz-3,4-dehydro-ʟ-proline **7a** was isolated by silica gel (96% $CH_2Cl_2$, 3% MeOH, 1% AcOH, $R_f$ = 0.25) for 0.120 g of total recovered product, a 72% yield. $^1$H-NMR (400 MHz, $CDCl_3$) δ 9.91 (b, 1H [COOH]), 7.37-7.27 (m, 5H), 5.98-5.93 (ddd, J = 21.3$_\Sigma$, 14.9, 6.11, 2.0 Hz 1H), 5.82-5.75 (ddd, J = 35.7$_\Sigma$, 29.6, 6.1, 2.2 Hz, 1H), 5.25-5.13 (m, J = 1.7 Hz, 2H), 5.06 (dd, J = 4.0$_\Sigma$, 2.2 Hz, 1H), 4.32-4.27 (m, J = 2.2, 2.0, 1.7 Hz, 2H).

*N-Cbz-3,4-dehydro-ʟ-proline benzyl ester* **7**. An open flask containing a mixture of 0.551 g (1.28 mmol) of N-Cbz-*trans*-4-methylxanthate-ʟ-proline benzyl ester **5a**, 1 mL ethanol, 10 mL water, and 68 mg $Na_2CO_3$ (anhydrous; 0.5 eq; 0.64 mmol) was placed into a microwave on medium low (100 watt noncontinous irradiation, retail purchased Sunbeam microwave) for ~10 min. The contents were allowed to cool to rt (keep in hood). The reaction mixture was diluted with 20 mL water and then extracted with 3 x 50 mL $Et_2O$, washed 2 x 50 mL water, dried over $Na_2SO_4$, and concentrated to a crude orange/brown colored oil. The product N-Cbz-3,4-dehydro-ʟ-proline benzyl ester **7** was isolated by silica gel (70% hexanes, 30% ethyl acetate, $R_f$ = 0.25) for 0.362 g of total recovered product, an 84% yield. $^1$H-NMR (400 MHz, $CDCl_3$) δ 7.40-7.25 (m, 10H), 6.01-5.94 (ddd, J = 19.0$_\Sigma$, 5.9, 1.5 Hz, 1H), 5.80-5.74 (ddd, J = 16.8$_\Sigma$, 5.9, 2.2 Hz, 1H), 5.26-5.04 (m, 5H), 4.40-4.26 (m, J = 5.9$_\Sigma$, 2.2, 1.5 Hz, 2H).

*N-Cbz-trans-3-hydroxy-cis-4-methoxy-ʟ-proline benzyl ester* **13a.** To 0.514 g (1.45 mmol) of N-Cbz-*trans*-3,4-epoxy-ʟ-proline benzyl ester **8a** was added 5 mL of dry $CH_2Cl_2$ (anhyrous $MgSO_4$

treated) and 2.0 mL anhydrous MeOH (~34 eq.; 50 mmol). The flask was capped, purged with argon and chilled on an ice bath with argon balloon. Once cooled, 1.15 mL (48%; 3.0 eq.; 4.36 mmol) of $BF_3$•$Et_2O$ was added to the mixture. The mixture was allowed to return to rt and stir over night. The reaction mixture was quenched by the addition of 20 mL saturated $NaHCO_3$ solution which was allowed to stir for ~30 min. The mixture was then diluted with 100 mL $CH_2Cl_2$, washed with 3 x 100 mL water, 50 mL brine, dried over $Na_2SO_4$ and concentrated to a solid foam. The crude containing N-Cbz-*trans*-3-hydroxy-*cis*-4-methoxy-L-proline benzyl ester, 467 mg, and unreacted epoxide starting material, 115 mg, were isolated by silica gel (75% hexanes, 25% ethyl acetate, $R_f$ product = 0.1 ; $R_f$ epoxide = 0.3) for a total yield of 83% intermediate product and complete recovery in molecular equivalents of unreacted epoxide starting material. The isolated product **13a** was used in the next step without further purification. $^1$H-NMR (500 MHz, $CDCl_3$) (2 conformational isomers) δ 7.33-7.25 (m, 10H), 5.27-5.01 (m, 4H), 4.50 and 4.40 (s, 1H), 3.85-3.75 (m, J = 4.89, 4.4 Hz, 2H), 3.70 (d, J = 4.9 Hz, 1H), 3.57 (m, 1H), 3.15 and 3.12 (s, 3H).

*trans-3-hydroxy-cis-4-methoxy-L-proline* **14a** (CSE115). To 0.417 g (1.08 mmol) of Cbz-3-*trans*-hydroxy-4-*cis*-methoxy-L-proline benzyl ester was added 4 mL methanol which was then transferred to a pressure flask containing 42 mg of 10% Pd/carbon (0.1 eq. weight), 1 mL water and 42 mg (~0.5 eq.) $NH_4OAc$. The pressure flask was connected to a Parr shaker, the air was removed by vacuum and replaced with 40 psi $H_2$ (g). The assembly was allowed to shake for 3 h. The pressure flask was vented and the solution diluted with 10 mL water and then filtered through CM-cellulose (carboxy-methyl-cellulose), using water for consecutive washes. The filtrate was then frozen and placed onto a lyophilizer until dry. The remaining residue was washed 3 x 10 mL ethyl acetate / $CH_2Cl_2$ (1:1), and dried leaving 0.172 g of a white solid, *trans*-3-

hydroxy-*cis*-4-methoxy-L-proline **14a**. $^1$H-NMR (500 MHz, $D_2O$ 4.75) δ 4.65 (s, 1H, H-2), 4.02 (s, 1H, H-1), 3.91 (s, 1H, H-3), 3.53 (s, 2H, H-4 and 5), 3.25 (s, 3H). $^{13}$C-NMR (500 MHz, $D_2O$) δ 170.7, 82.8, 74.8, 67.4, 56.3, 48.9. HRMS *m/e* calcd. For $C_6H_{12}NO_4$ = 162.0749, found 162.0766. $[\alpha]^{22}_{Na}$ = -15° (c = 0.14, EtOH).

*N-Cbz-cis-3-hydroxy-trans-4-methoxy-L-proline benzyl ester and N-Cbz-trans-3-methoxy-cis-4-hydroxy-L-proline benzyl ester* **15a** and **16a.** To 0.280 g (0.792 mmol) of N-Cbz-*cis*-3,4-epoxy-L-proline benzyl ester **8b** was added 3 mL dry $CH_2Cl_2$ (anhydrous $MgSO_4$ treated) and 1 mL anhydrous MeOH (~30 eq.; 25 mmol). The flask was purged with argon and cooled on ice bath for 10 min. Once cooled, 0.625 mL of $BF_3 \bullet Et_2O$ (48%; 3.0 eq.; 2.38 mmol) was added dropwise and the solution was allowed to return to rt. After stirring overnight the reaction was quenched by the addition of 20 mL saturated solution of $NaCO_3$. This was then further diluted with 100 mL $CH_2Cl_2$ and washed 3 x 100 mL water and 1 x 100 mL brine, dried with $Na_2SO_4$ and filtered. The crude containing the two regioisomer products N-Cbz-*cis*-3-hydroxy-*trans*-4-methoxy-L-proline benzyl ester and N-Cbz-*trans*-3-methoxy-*cis*-4-hydroxy-L-proline benzyl ester, 0.288 g, were isolated together by silica gel (70 % hexanes, 30% ethyl acetate, $R_f$ = 0.15) a 94% yield. The intermediates were used in the next step without further purification.

*N-Cbz-cis-3-hydroxy-4-trans-methoxy-L-proline*. To 0.410 g (1.06 mmol) mixture of N-Cbz-*cis*-3-hydroxy-*trans*-4-methoxy-L-proline benzyl ester **15a** and N-Cbz-*trans*-3-methoxy-*cis*-4-hydroxy-L-proline benzyl ester **16a** was added 10 mL THF, 1 mL water and 72 mg (1.2 eq.; 1.28 mmol) KOH. The mixture was allowed to stir for 2 h at rt where it was observed by TLC (hexanes : EtOAc, 7 : 3) that all starting material was consumed. The mixture was concentrated to an oil and diluted with 20 mL water. The pH was adjusted to ~2 by addition of $H_3PO_4$ and NaCl was added until saturated. The mixture was then extracted with 2 x 100 mL ethyl acetate. The separated organic layers were combined, washed 20 mL pH of ~2 brine, dried with $NaSO_4$, filtered and concentrated. The crude mixture, 290 mg (0.979 mmol; 92% yield), was separated on silica gel (94% $CH_2Cl_2$, 5% MeOH, 1% AcOH, $R_f$ = 0.2) and again concentrated. A portion of this regioisomer mixture was isolated by reverse phase HPLC (C18, 250mm x 21.2mm, 10µ) (88% 0.05 M $NH_4OAc$, 12% acetonitrile, 9 mL/min) at retention times of 44 min. for N-Cbz-3-*cis*-hydroxy-4-*trans*-methoxy-ʟ-proline or 46 min for the 4-*cis*-hydroxy regio isomer and concentrated by freezing / lyophilization. The isolated free acid N-Cbz-3-*cis*-hydroxy intermediate of **15a** was used in the next step without further purification. $^1$H-NMR (500 MHz, $D_2O$ 4.65) (2 conformational isomers) δ 7.26-7.20 (m, 5H), 5.02-4.95 (dd, J = 25.4, 12.2 Hz, 1H), 4.94 (s, 1H), 4.34 and 4.33 (s, 1H), 4.23 and 4.18 (d, J = 6.4 Hz, 1H), 3.74 and 3.73 (s, 1H), 3.74-3.73 (m, 1H), 3.50 and 3.42 (d, 12.0 Hz, 1H), 3.23 and 3.21 (s, 3H).

*cis-3-hydroxy-trans-4-methoxy-ʟ-proline* **17a** (CSE124). To 90.0 mg (0.305 mmol) of N-Cbz-*cis*-3-hydroxy-4-*trans*-methoxy-ʟ-proline was added 3 mL MeOH and 1 mL water. This was transferred to a pressure flask containing 10 mg of wet 10% Pd/carbon (0.1 eq. weight) and 12 mg $NH_4OAc$ (0.5 eq.). The reaction flask was placed onto a Parr shaker, the air was removed by vacuum and replaced with $H_2$ (g) to 40 psi. The flask was allowed to shake for 8 h where the flask was vented and mixture was diluted with ~ 20 mL water. The contents were then filtered through CM-cellulose using water in consecutive washes. The filtrate was frozen and lyophilized to a white residue and then rinsed with 3 x 20 mL ethyl acetate / dichloromethane (1:1) and dried leaving a white solid residue; 44.0 mg of *cis*-3-hydroxy-*trans*-4-methoxy-ʟ-proline **17a** a 90% yield. $^1$H-

NMR (500 MHz, $D_2O$ 4.75) δ 4.56 (d, J = 3.8 Hz, 1H, H-2), 4.22 (d, J = 3.8 Hz, 1H, H-1), 4.01 (d, J = 3.5 Hz, 1H, H-3), 3.62-3.59 (dd, J = 13.0, 3.5 Hz, 1H, H-5), 3.41-3.39 (d, J = 13.0 Hz, 1H, H-4), 3.35 (s, 3H). $^{13}$C-NMR (500 MHz, $D_2O$) δ 170.3, 83.9, 72.4, 65.2, 56.5, 48.36. HRMS *m/e* calcd. For $C_6H_{11}NO_4$ = 162.0749, found 162.0753. $[\alpha]^{22}_{Na}$ = -10° (c = 0.23, EtOH).

*N-Cbz-trans-3-hydroxy-cis-4-isopropoxy-L-proline benzyl ester* **13b**. To 0.460 g (1.30 mmol) of N-Cbz-*trans*-3,4-epoxy-L-proline benzyl ester **8a** was added 4 mL of dry ($MgSO_4$ treated) $CH_2Cl_2$ and 2.0 mL isopropanol (~20 eq.; 26 mmol). The flask was capped, purged with argon and chilled on an ice bath with argon balloon. Once cooled, 1.03 mL of $BF_3$•$Et_2O$ (48%; 3 eq.; 3.90 mmol) was added to the mixture. The mixture was allowed to return to rt and stir over night. The reaction mixture was quenched by the addition of 20 mL saturated $NaHCO_3$ solution which was allowed to stir for ~30 min. The mixture was then diluted with 100 mL $CH_2Cl_2$, washed with water (3 x 200 mL), washed with 50 mL brine, dried over $Na_2SO_4$ and concentrated. The final product N-Cbz-*trans*-3-hydroxy-*cis*-4-isopropoxy-L-proline benzyl ester **13b**, 430 mg, and unreacted epoxide starting material, were isolated by silica gel (hexanes : ethyl acetate 3 : 1, $R_f$ product = 0.15 ; $R_f$ epoxide = 0.3) for a total yield of 76% product and complete recovery in molecular equivalents of unreacted starting material (epoxide). The material was used in the next step without further purification. $^1$H-NMR (500 MHz, $CDCl_3$) (2 conformational isomers) δ 7.33-7.22 (m, 10H), 5.28-5.03 (m, 4H), 4.49 and 4.44 (s, 1H), 4.43 and 4.37 (s, 1H), 3.91-3.89 (d, J = 2.4 Hz, 1H), 3.86-3.82 and 3.81-3.77 (dd, J = 11.5, 4.9 Hz, 1H), 3.65-3.58 (m, J = 6.1 Hz, 1H), 3.53-3.50 (d, J = 2.4 Hz, 1H), 1.07-1.00 (m, J = 5.9 Hz, 6H). $^{13}$C-NMR (500 MHz, $CDCl_3$) (2 conformational isomers) δ 167.2 and 166.8, 152.9 and 152.4, 133.8 and 133.0, 125.9, 125.9, 125.8, 125.6, 125.5, 125.4, 125.4, 125.4,

125.3, 125.2, 76.5, 75.8, 74.8, 74.6, 74.6, 74.3, 67.8 and 67.7, 64.8 and 64.7, 64.4 and 64.3, 63.7 and 63.2, 48.2 and 48.0, 19.7 and 19.6, 19.5 and 19.5.

*trans-3-hydroxy-cis-4-isopropoxy-L-proline* **14b** (CSE122). To 0.190 g (0.460 mmol) of Cbz-3-*trans*-hydroxy-4-*cis*-methoxy-L-proline benzyl ester was added 4 mL methanol which was then transferred to a pressure flask containing 19 mg (0.1 eq. weight) of 10 % Pd / carbon, 1 mL water and 18 mg (0.5 eq.) of $NH_4OAc$. The pressure flask was connected to a Parr shaker, the air was removed by house vacuum and replaced with 40 psi $H_2$ (g). The assembly was allowed to shake for 3 h. The pressure flask was vented and the solution diluted with 10 mL water and then filtered through CM-cellulose (carboxy-methyl-cellulose), using excess water for consecutive washes. The filtrate was then frozen and placed onto a lyophilizer until dry. The remaining residue was washed 3 x 20 mL EtOAc / $CH_2Cl_2$ (1:1), and dried leaving 80.0 mg of a white solid, *trans*-3-hydroxy-*cis*-4-isopropoxy-L-proline **14b** with a yield of 92%. $^1$H-NMR (500 MHz, $D_2O$ 4.75) δ 4.59 (s, 1H, H-2), 4.10 (d, J = 4.0 Hz, 1H, H-3), 3.99 (s, 1H, H-1), 3.75-3.70 (m, J = 6.1 Hz, 1H), 3.56-3.53 (dd, J = 12.7, 4.0 Hz, 1H, H-4), 3.47-3.44 (d, J = 12.7 Hz, 1H, H-5), 1.06 (d, J = 6.1 Hz, 3H), 1.01 (d, J = 6.1 Hz, 3H). $^{13}$C-NMR (500 MHz, $D_2O$) δ 170.7, 78.4 and 78.2, 75.7 and 75.7, 70.9 and 70.8, 67.6 and 67.4, 49.7, 21.4 and 21.4, 20.6 and 20.5. HRMS *m/e* calcd. For $C_8H_{16}NO_4$ = 190.1079, found 190.1089. $[\alpha]^{22}_{Na}$ = -5° (c = 0.24, EtOH).

*N-Cbz-cis-3-hydroxy-trans-4-isopropoxy-L-proline benzyl ester and N-Cbz-trans-3-isopropoxy-cis-4-hydroxy-L-proline benzyl ester* ***15b*** *and* ***16b***. To 0.407 g (1.15 mmol) of N-Cbz-*cis*-3,4-epoxy-L-proline benzyl ester **8b** was added 5 mL (dry $MgSO_4$ treated) $CH_2Cl_2$ and 2 mL isopropanol (~23 eq.; 26 mmol). The flask was purged with argon and cooled on ice bath for 10 min. Once cooled,

0.910 mL of $BF_3$•$Et_2O$ (48%; 3 eq.; 3.45 mmol) was added dropwise and the solution was allowed to return to rt. After stirring overnight the reaction was quenched by the addition of 20 mL saturated solution of $NaCO_3$. This was then further diluted with 20 mL $CH_2Cl_2$ and washed 3 x 100 mL water and 1 x 100 mL brine, dried with $Na_2SO_4$ and concentrated to an oil. The crude containing the two regioisomer products N-Cbz-*cis*-3-hydroxy-*trans*-4-isopropoxy-L-proline benzyl ester **15b** and N-Cbz-*trans*-3-isopropoxy-*cis*-4-hydroxy-L-proline benzyl ester **16b**, 0.440 g, were isolated together by silica gel (3:1 hexanes : EtOAc, $R_f$ = 0.20) a 92% yield. The material as a regioisomer mixture was used in the next step without further purification.

*N-Cbz-cis-3-hydroxy-4-trans-isopropoxy-L-proline*. To 0.400 g (0.967 mmol) of N-Cbz-*cis*-3-hydroxy-*trans*-4-isopropoxy-L-proline benzyl ester **15b** and N-Cbz-*trans*-3-isopropoxy-*cis*-4-hydroxy-L-proline benzyl ester **16b** was added 10 mL THF, 1 mL water and 81 mg KOH (1.5 eq.; 1.45 mmol). The mixture was allowed to stir for 2 h at rt until complete, as determined by TLC (50% hexanes, 49% ethyl acetate, 1% AcOH). The mixture was concentrated to an oil by rotovap and diluted with 20 mL water. The pH of solution was adjusted to ~2 by addition of $H_3PO_4$ and NaCl was added until saturated. The mixture was then extracted with 3 x 100 mL EtOAc. The separated organic layers were combined, dried with $Na_2SO_4$, filtered and concentrated to a crude residue. The free acid products, 280 mg (0.866 mmol; 89% yield), were isolated as a mixture by silica gel (60% hexanes, 39% ethyl acetate, 1% AcOH, $R_f$ = 0.1) and again concentrated. A portion of this regioisomer mixture was isolated by reverse phase HPLC (C18, 250mm x 21.2mm, 10µ) (85% 0.05 M $NH_4OAc$, 15% acetonitrile, 9 mL/min) at retention times of 48 min for N-Cbz-*cis*-3-hydroxy-4-*trans*-isopropoxy-L-proline & 52 min. for N-Cbz-*trans*-3-isopropoxy-*cis*-4-hydroxy-L-proline and stored in the freezer. The free acid derivative of **15b**, only, was concentrated by lyophilization and used in the next step without further purification.

*cis-3-hydroxy-trans-4-isopropoxy-L-proline, **17b*** (CSE127). To 21.0 mg (0.0650 mmol) of N-Cbz-*cis*-3-hydroxy-4-*trans*-isopropoxy-L-proline was added 3 mL MeOH and 1 mL water. This was transferred to a pressure flask containing 5 mg of wet 10% (0.25 eq. weight) Pd/carbon and 5 mg (0.5 eq) $NH_4OAc$. The reaction flask was placed onto a Parr shaker, the air was removed by vacuum and replaced by 40 psi $H_2$ (g). The flask was allowed to shake for 8 h where the flask was vented and mixture was diluted with ~ 20 mL water. The contents were then filtered through CM-cellulose using excess water for consecutive washes. The filtrate was frozen and lyophilized to a white solid and washed with EtOAc / $CH_2Cl_2$ 1:1 (3 x 20 mL) and dried leaving a white solid; 11.6 mg of *cis*-3-hydroxy-*trans*-4-isopropoxy-L-proline a 94% yield. $^1$H-NMR (500 MHz, $D_2O$ 4.75) δ 4.48 (d, J = 3.9 Hz, 1H, H-2), 4.24 (d, J = 3.9 Hz, 1H, H-1), 4.21 (d, J = 3.9 Hz, 1H, H-3), 3.82-3.78 (m, J = 5.9, 3.9 Hz1H), 3.63 (dd, J = 12.7, 3.9 Hz, 1H, H-5), 3.32 (d, J = 12.7 Hz, 1H, H-4), 1.12-1.10 (m, J = 5.9, 3.9 Hz, 6H). $^{13}$C-NMR (500 MHz, $D_2O$) (2 conformational isomers) δ 79.9, 73.5, 71.4, 65.3, 49.3, 21.3, 21.2. HRMS *m/e* calcd. For $C_8H_{16}NO_4$ = 190.1079, found 190.1092. $[\alpha]^{22}_{Na}$ = -7 ° (c = 0.13, EtOH).

*N-Cbz-trans-3-hydroxy-cis-4-benzyloxy-L-proline benzyl ester **13c***. To 0.564 g (1.59 mmol) of N-Cbz-*trans*-3,4-epoxy-L-proline benzyl ester **8a** was added 5 mL of dry ($MgSO_4$ treated) $CH_2Cl_2$ and 2.5 mL anhydrous benzyl alcohol (~15 eq; 24 mmol). The flask was capped, purged with argon and chilled on an ice bath under argon balloon. Once cooled, 1.26 mL (48%; 3 eq.; 4.78 mmol) of $BF_3 \bullet Et_2O$ was added to the mixture. The mixture was allowed to return to rt and stir for 48 h.

The reaction was quenched by the addition of 40 mL saturated $NaHCO_3$ solution and allowed to stir for 30 min. The mixture was then diluted with 100 mL $CH_2Cl_2$, washed water (2 x 200 mL), washed with 50 mL brine, dried over $Na_2SO_4$ and concentrated to a white solid. The final product N-Cbz-*trans*-3-hydroxy-*cis*-4-benzyloxy-ʟ-proline benzyl ester, 0.710 mg (1.54 mmol), was isolated by silica gel (85% hexanes, 15% ethyl acetate, $R_f$ = 0.10) for a total yield of 96%. The intermediate was used in the next synthetic step without further purification. $^1$H-NMR (500 MHz, $CDCl_3$) (2 conformational isomers) δ 7.33-7.18 (m, 15H), 5.18-5.10 (m, 1H), 5.05 (s, 1H), 5.03-4.92 (m, 1H), 4.55 (s, 1H), 4.50-4.39 (m, 2H), 3.91 (dd, J = 10.3, 5.1 Hz, 1H), 3.85-3.77 (m, J = 11.7, 5.1 Hz, 1H), 3.67-3.62 (dd, J = 12.0, 11.7 Hz, 1H), 3.42 (s, 2H).

*trans-3-hydroxy-cis-4-benzyloxy-ʟ-proline,* ***14c*** (CSE121). To 0.710 g (1.56 mmol) of Cbz-3-*trans*-hydroxy-4-*cis*-benzyloxy-ʟ-proline benzyl ester **13c** was added 5 mL methanol which was then transferred to a pressure flask containing 71 mg (0.1 eq. weight) of 10% Pd/carbon, 0.3 mL water and 59.3 mg (0.5 eq.; 0.769 mmol) $NH_4OAc$. The pressure flask was connected to a Parr shaker, the air was removed by vacuum and replaced with 40 psi $H_2$ (g). The assembly was allowed to shake for 3 h. The pressure flask was vented and the contents were diluted with 30 mL water and filtered through CM-cellulose using excess water for consecutive washes. The filtrate was then frozen and placed onto a lyophilizer until dry. The remaining residue was washed EtOAc / $CH_2Cl_2$ 1:1 (3 x 20 mL), and dried leaving 0.340 mg of a white crude residue *trans*-3-hydroxy-*cis*-4-benzyloxy-ʟ-proline, a yield of 93%. A portion of this crude product was isolated by reverse phase HPLC (C18, 250mm x 21.2mm, 10μ) (92% 0.05 M $NH_4OAc$, 8% acetonitrile, 9 mL/min) at a retention time of 42 min. The collected fractions containing the desired product were concentrated by lyophilization. Once dry ~20 mL of water was added, the solution was again frozen and lyophilized until dry removing any remaining $NH_4OAc$. This was repeated until $NH_4OAc$ was no longer present, leaving a white solid *trans*-3-hydroxy-*cis*-4-benzyloxy-ʟ-proline, **14c** (CSE 121). $^{1}$H-NMR (500 MHz, $D_2O$ 4.75) δ 7.37-7.30 (m, 5H), 4.71 (s, 1H, H-2), 4.56-4.47 (dd, J = 31.1, 11.5 Hz, 2H, methylene), 4.09 (s, 1H, H-3), 4.04 (s, 1H, H-1), 3.55-3.54 (d, J = 1.5 Hz, 2H, H-4 and 5). $^{13}$C-NMR (500 MHz, $D_2O$) δ 170.5, 136.7, 128.6, 128.2, 128.2, 80.8, 75.1, 71.0, 67.5, 49.4. HRMS *m/e* calcd. For $C_{12}H_{16}NO_4$ = 238.1079, found 238.1094. $[\alpha]^{22}_{Na}$ = -13° (c = 0.08, EtOH).

*N-Cbz-cis-3-hydroxy-trans-4-benzyloxy-ʟ-proline benzyl ester* ***15c*** *and N-Cbz-trans-3-benzyloxy-cis-4-hydroxy-ʟ-proline benzyl ester* ***16c***. To 0.484 g (1.37 mmol) of N-Cbz-*cis*-3,4-epoxy-ʟ-proline benzyl ester **8b** was added 5 mL of dry ($MgSO_4$ treated) $CH_2Cl_2$, 20 mL benzyl alcohol (treated with molecular civs), and 13.0 mg (0.10 eq) of p-toluensulfonic acid monohydrate. The mixture was heated to ~60°C and stirred overnight until the reaction appeared to be complete as determined by TLC (hexanes : ethyl acetate, 1:1, $R_f$ = 0.4). The solution was then allowed to cool to rt, diluted with 100 mL $Et_2O$ and washed with water (3 x 100 mL). After drying with $Na_2SO_4$, the separated organic phase was filtered and run through a silica plug using 90% hexanes 10% ethyl acetate to remove most of the excess benzyl alcohol. The product was then eluted with 1:1 hexanes / ethyl acetate and concentrated. The final products N-Cbz-*cis*-3-hydroxy-*trans*-4-benzyloxy-ʟ-proline benzyl ester and N-Cbz-*trans*-3-benzyloxy-*cis*-4-hydroxy-ʟ-proline benzyl ester, 0.426 g (0.923 mmol), were isolated by silica gel (7: 3 hexanes : ethyl acetate, $R_f$ product = 0.15) for a total yield of 67% including both regioisomers. Note: this reaction was repeated using instead 3.0 eq. of $BF_3$•$Et_2O$ catalyst instead of tosylic acid and under similar conditions as with the *trans*-epoxide – benzyl alcohol addition and yields improved dramatically, 197 mg of the two regioisomers were isolated starting from 160 mg of starting *cis*-epoxide intermediate. The resulting products as a regioisomer mixture were used in the next step without further purification.

*cis-3-hydroxy-trans-4-benzyloxy-L-proline **17c*** (CSE113). To 0.426 g (0.923 mmol) of Cbz-*cis*-hydroxy-benzyloxy-L-proline benzyl ester intermediates, **15c** and **16c**, was added 5 mL methanol which was then transferred to a pressure flask containing 40 mg (0.1 eq. weight) of 10% Pd/carbon, 2 mL MeOH, 1 mL water and 36 mg (~0.5 eq.; 0.47 mmol) of $NH_4OAc$. The pressure flask was connected to a Parr shaker, the air was removed by vacuum and replaced with 40 psi $H_2$ (g). The assembly was allowed to shake for 1.5 h where the reaction was complete as determined by TLC (7 : 3 hexanes/ EtOAc). The pressure flask was vented and the contents were diluted with 30 mL water and filtered through CM-cellulose (carboxy-methyl-cellulose), using excess water for consecutive washes. The filtrate was then frozen and lyophilized until dry. The remaining solid was washed with EtOAc / $CH_2Cl_2$ 1:1 (3 x 20 mL), and dried leaving 0.206 mg of a white solid, a mixture of *cis*-3-hydroxy-*trans*-4-benzyloxy-L-proline **17c** and *trans*-3-benzyloxy-*cis*-4-hydroxy-L-proline **18c**. A portion of this crude regioisomer mixture was isolated by reverse phase HPLC (C18, 250mm x 21.2mm, 10µ) (92% 0.05 M $NH_4OAc$, 8% acetonitrile, 9 mL/min) collecting the *cis*-3-hydroxy prolinol product **17c** which were concentrated via lyophilization. Fractions containing the 4-hydroxy prolinol **18c** were frozen and saved. Once dry ~20 mL of water was added, the solution was again frozen and lyophilized until dry. This was repeated until all remaining $NH_4OAc$ was removed, leaving *cis*-3-hydroxy-*trans*-4-benzyloxy-L-proline, **17c** (CSE113). $^1$H-NMR (500 MHz, $D_2O$ 4.75) δ 7.38-7.35 (m, 5H), 4.60 (s, 1H, H-2), 4.59 (s, 2H, methylene), 4.27-4.26 (d, J = 3.9 Hz, 1H, H-1), 4.21-4.20 (d, J = 3.9 Hz, 1H, H-3), 3.66-3.62 (dd, J =

13.2, 3.9 Hz, 1H, H-5), 3.44-3.42 (d, J = 13.2 Hz, 1H, H-4). $^{13}$C-NMR (500 MHz, $D_2O$) δ 172.8, 139.2, 131.3, 131.1, 130.9, 84.7, 75.4, 73.8, 67.9, 51.4. HRMS *m/e* calcd. For $C_{12}H_{16}NO_4$ = 238.1079, found 238.1064. $[\alpha]^{22}_{Na}$ = -20° (c = 0.22, EtOH).

*(2S, 3R, 4R) N-Cbz-trans-3-hydroxy-cis-4-methyloxy-1-napthyl-L-proline benzyl ester* **13d**. To 0.285 g (0.806 mmol) of N-Cbz-*trans*-3,4-epoxy-L-proline benzyl ester **8a** was added 2 mL of dry ($MgSO_4$ treated) $CH_2Cl_2$ and 0.191 g (98%, 1.5 eq.; 1.20 mmol) of 1-naphthylenemethanol. The flask was capped, purged with argon and chilled on an ice bath with argon balloon. Once cooled, 64 µL (48%; 0.3 eq.; 0.242 mmol) of $BF_3$•$Et_2O$ was added to the stirring solution mixture. The mixture was allowed to return to rt and stir over night. The reaction was quenched by the addition of 20 mL saturated $NaHCO_3$ solution and allowed to stir for ~30 min. The contents of the flask were then diluted with 100 mL $CH_2Cl_2$, washed with water (3 x 100 mL), 50 mL brine, dried over $Na_2SO_4$ and concentrated to a white crude. The final product (2S, 3R, 4R) N-Cbz- *trans*-3-hydroxy-*cis*-4-methyloxy-1-napthyl-proline benzyl ester **13d**, 0.220 mg (0.430 mmol), was isolated by silica gel (7 : 3 hexanes, EtOAc, $R_f$ = 0.1) a yield of 53%, only a portion of product was isolated, and recovery of 30 mg epoxide starting material. The product was used in the next synthetic step without further purification.

*(2S, 3R, 4R) trans-3-hydroxy-cis-4-methyloxy-1-napthyl-L-proline* **14d** (CSE132). To 0.120 g (0.235 mmol) of (2S, 3R, 4R)-N-Cbz-3-hydroxy-4-o-methyl-α-napthyl-proline benzyl ester **13d** was added 5 mL methanol which was heated to dissolve and then transferred to a pressure flask containing 20 mg of 10% Pd/carbon (0.167 eq. weight), and 0.5 mL water. The pressure flask was connected to a Parr shaker, the air was removed by vacuum and replaced with $H_2$ (g) to 40 psi. The assembly

was allowed to shake over night until it was clear by TLC that all the starting material was consumed (50/49/1; hexanes / EtOAc / AcOH). The pressure flask was vented and the contents were diluted with 30 mL water, followed by filtration through CM-cellulose (carboxy-methyl-cellulose), using excess water for consecutive washes. The filtrate was then frozen and placed onto a lyophilizer until dry. The remaining residue was washed EtOAc / $CH_2Cl_2$ 1:1 (3 x 20 mL), and dried leaving 58.0 mg of a white solid, primarily consisting of the desired prolinol product, a yield of 86%. A portion of this prolinol product crude was isolated by reverse phase HPLC (C18, 250mm x 21.2mm, 10μ) (75% 0.05 M $NH_4OAc$ (aq), 25% acetonitrile, 9 mL/min) at a retention time of ~44 min. The collected fractions containing the desired prolinol were combined and concentrated by lyophilization. Once dry, 20 mL of water was added, the solution was again frozen and lyophilized until dry. This was repeated until $NH_4OAc$ was no longer present, leaving a white solid (2S, 3R, 4R) 3-hydroxy-4-o-methyl-α-napthyl-proline, **14d** (CSE132). $^1$H-NMR (500 MHz, $d_6$-DMSO) δ 8.84 (b, 2H ($-NH_2^+$), 8.05 (d, J = 7.8 Hz, 1H), 7.93 (d, J = 7.3 Hz, 1 H), 7.87 (d, J = 7.8 Hz, 1H), 7.57-7.51 (m, 3H), 7.46 (dd, J = 7.8, 7.3 Hz, 1H), 5.66 (b, 1H, (-OH)), 4.95-4.88 (dd, J = 19.6, 12.2 Hz, 2H), 4.58 (s, 1H, H-2), 3.94 (d, J = 2.4 Hz, 1H, H-3), 3.50 (s, 1H, H-1), 3.47 (d, J = 12.7 Hz, 1H, H-5), 3.28-3.25 (dd, J = 12.7, 3.4 Hz, 1H, H-4). $^{13}$C-NMR (500 MHz, $d_6$-DMSO) δ 166.5, 133.3, 131.0, 128.3, 128.2, 126.3, 125.8, 125.3, 124.0, 119.7, 113.6, 82.1, 75.8, 69.7, 68.2, 48.6. HRMS *m/e* calcd. For $C_{16}H_{18}NO_4$ = 288.1236, found 288.1230. $[\alpha]^{22}_{Na}$ = -18° (c = 0.1, DMSO).

*(2S, 3R, 4R) N-Cbz-trans-3-hydroxy-cis-4-methyloxy-p-biphenyl-L-proline benzyl ester* **13e**. To 0.305 g (0.862 mmol) of N-Cbz-*trans*-3,4-epoxy-L-proline benzyl ester **8a** was added 2 mL of dry ($MgSO_4$ treated) $CH_2Cl_2$ and 0.318 g (98%, 2.0 eq.; 1.72 mmol) of 4-biphenylmethanol. The flask was capped, evacuated via vacuum line and chilled on an ice bath under argon balloon. Once

cooled, 91 µL (48%; 0.4 eq.; 0.345 mmol) of $BF_3$•$Et_2O$ was added to the stirring solution. The mixture was allowed to return to rt and stir over night. The reaction was quenched by the addition of 20 mL saturated $NaHCO_3$ (aq) and allowed to stir for an additional 30 min. The contents of the flask were then diluted with 100 mL $CH_2Cl_2$, washed with water (3 x 100 mL), 50 mL brine, dried over $Na_2SO_4$ and concentrated to a white solid. The final product (2S, 3R, 4R) N-Cbz-*trans*-3-hydroxy-*cis*-4-methyloxy-p-biphenyl-L-proline benzyl ester **13e**, 0.395 mg (0.735 mmol), was isolated by silica gel (7 : 3 hexanes : EtOAc, $R_f$ = 0.1), a yield of 85%. Some remaining epoxide starting material was recovered. The product was used in the next step without further purification.

*(2S, 3R, 4R) trans-3-hydroxy-cis-4-methyloxy-p-biphenyl-L-proline* ***14e*** (CSE131). To 0.105 g (0.195 mmol) of (2S, 3R, 4R)-N-Cbz-3-hydroxy-4-o-methyl-p-biphenyl-proline benzyl ester was added 5 mL methanol which was heated to dissolve and then transferred to a pressure flask containing 11 mg of 10% Pd/carbon (0.10 eq. weight), 8 mg (0.5 eq) $NH_4OAc$ and 0.5 mL water. The pressure flask was connected to a Parr shaker, the air was removed by vacuum and replaced with 40 psi $H_2$ (g). The assembly was allowed to shake for 2 h until the reaction had completed as determined by TLC (50/49/1; hexanes/ethyl acetate/AcOH). The pressure flask was vented and the contents were diluted with 30 mL water, followed by filtration through CM-cellulose using excess water for consecutive washes. The filtrate was then frozen and lyophilized until dry. The remaining residue was washed EtOAc / $CH_2Cl_2$ 1 : 1 ( 3 x 20 mL), and dried over $Na_2SO_4$ leaving 51.0 mg of a white solid crude, primarily consisting of the desired prolinol product, a yield of 84%. A portion of this crude product was isolated by reverse phase HPLC (C18, 250mm x 21.2mm, 10µ) (75% 0.05 M $NH_4OAc$, 25% acetonitrile, 9 mL/min) at a retention time of ~38 min. The collected

fractions containing the desired prolinol were combined and concentrated by lyophilization. Once dry, 20 mL of water was added, the solution was again frozen and lyophilized until dry (removes remaining $NH_4OAc$). This was repeated until $NH_4OAc$ was no longer present, leaving a white residue 2S, 3R, 4R) *trans*-3-hydroxy-*cis*-4-methyloxy-p-biphenyl-ʟ-proline **14e** (CSE131). $^1$H-NMR (500 MHz, $d_6$-DMSO 2.50) δ 8.82 (b, 1.5H, $NH_3^+$), 7.66-7.65 (d, J = 7.6 Hz, 2H), 7.62-7.60 (d, J = 7.8 Hz, 2H), 7.47-7.44 (dd, J = 7.6, 7.3 Hz, 2H), 7.41-7.40 (d, J = 7.6 Hz, 2H), 7.37-7.34 (dd, J = 7.3, 7.1 Hz, 1H), 5.64 (s, 1H, OH), 4.54 (s, 1H, H-2), 4.53-4.46 (dd, J = 21.5, 22.7 Hz, 2H, methylene), 3.86 (s, 1H, H-3), 3.49 (s, 1H, H-1), 3.46-3.43 (d, J = 12.5 Hz, 1H, H-5), 3.33 ($H_2O$), 3.27-3.25 (d, J = 10.5 Hz, 1H, H-4). $^{13}$C-NMR (500 MHz, $d_6$-DMSO) δ 170.2, 140.0, 139.2, 137.1, 128.9, 128.3, 127.4, 126.6, 126.4, 81.8, 75.4, 69.5, 68.2, 46.7. HRMS *m/e* calcd. For $C_{18}H_{20}NO_4$ = 314.1392, found 314.1378. $[\alpha]^{22}_{Na}$ = -7° (c = 0.12, DMSO).

*(2S, 3R, 4R) N-Cbz-trans-3-hydroxy-cis-4-methyloxy-p-diphenylether-proline benzyl ester* **13f**. To 0.275 g (0.778 mmol) of N-Cbz-*trans*-3,4-epoxy-ʟ-proline benzyl ester **8a** was added 3 mL of dry ($MgSO_4$ treated) $CH_2Cl_2$ and 1 mL (7.34 eq.; 5.74 mmol) of 3-phenoxybenzyl alcohol. The flask was capped, purged with argon and chilled on an ice bath under argon balloon. Once cooled, 0.614 mL (48%; 3 eq.; 2.33 mmol) of $BF_3 \bullet Et_2O$ was added to the mixture. The mixture was allowed to return to rt and stir for 48 h. The reaction was quenched by the addition of 20 mL saturated $NaHCO_3$ and allowed to stir for 30 min. The mixture was then diluted with 60 mL $CH_2Cl_2$, washed with water (3 x 200 mL), washed with 50 mL brine, dried over $Na_2SO_4$ and concentrated via rotovap to a solid residue. The final product **(**2S, 3R, 4R) N-Cbz-*trans*-3-hydroxy-*cis*-4-methyloxy-p-diphenylether-proline benzyl ester **13f**, 0.270 g (0.488 mmol), was isolated by

silica gel (7 : 3 hexanes : EtOAc, $R_f$ = 0.10) for a total yield of 63%. The product was used in the next step without further purification.

*(2S, 3R, 4R) trans-3-hydroxy-cis-4-methyloxy-p-diphenylether-L-proline* **14f** (CSE130). To 0.270 g (0.488 mmol) of **(**2S, 3R, 4R) N-Cbz-*trans*-3-hydroxy-*cis*-4-methyloxy-p-diphenylether-proline benzyl **13f** ester was added 5 mL methanol which was then transferred to a pressure flask containing 30 mg of 10% Pd/carbon (0.1 eq. weight), 0.3 mL water and 19.0 mg (0.5 eq.; 0.246 mmol) $NH_4OAc$. The pressure flask was connected to a Parr shaker, the air was removed by vacuum and replaced with 40 psi $H_2$ (g). The assembly was allowed to shake for 2 h. The pressure flask was vented and the contents were diluted with 20 mL water, followed by filtration through CM-cellulose using excess water for consecutive washes. The filtrate was then frozen and placed onto a lyophilizer until dry. The remaining residue was washed with 1:1 EtOAc : $CH_2Cl_2$ (3 x 20 mL), and dried leaving 0.144 mg of a white solid crude, primarily consisting of the desired prolinol, a yield of 90%. A portion of this crude product was isolated by reverse phase HPLC (C18, 250mm x 21.2mm, 10µ) (75% 0.05 M $NH_4OAc$, 25% acetonitrile, 9 mL/min) at a retention time of ~42 min. The collected fractions containing the prolinol product were combined and concentrated by lyophilization. Once dry 20 mL of water was added, the solution was again frozen and lyophilized until dry (removes remaining $NH_4OAc$). This was repeated until $NH_4OAc$ was no longer present, leaving a white residue **(**2S, 3R, 4R) *trans*-3-hydroxy-*cis*-4-methyloxy-p-diphenylether-L-proline **14f** (CSE130). $^1$H-NMR (500 MHz, $d_6$-DMSO 2.50) δ 8.81 (b, 1.5H, $NH_3^+$), 7.40-7.37 (dd, J = $14.7_{\Sigma}$, 7.3 Hz, 2H), 7.34-7.31 (dd, J = 7.8, 7.6 Hz, 1H), 7.15-7.09 (m, J = 18.1, 7.6 Hz, 2H), 6.99 (m, 3H), 6.87 (d, J = 7.6, 1H), 5.62 (s, 1H, OH), 4.51 (s, 1H, H-2), 4.48-4.41 (dd, J = 23.0, 12.0 Hz, 2H, methylene), 3.83 (s, 1H, H-3), 3.47 (s, 1H, H-1), 3.43-3.41 (d, J = 12.2 Hz, 1H, H-5), 3.25-3.23 (d, J

= 11.0 Hz, 1H, H-4). $^{13}$C-NMR (500 MHz, $d_6$-DMSO) δ 169.1, 156.7, 156.4, 140.3, 130.1, 129.7, 123.4, 122.7, 118.5, 117.8, 117.5, 81.9, 75.4, 69.3, 68.1, 48.7. HRMS *m/e* calcd. For $C_{18}H_{20}NO_5$ = 330.1328, found 330.1341. $[\alpha]^{22}_{Na}$ = -29° (c = 0.14, DMSO).

*N-Cbz-3,4-dehydro-L-proline* **7a**. To 8.26 g (24.5 mmol) of N-Cbz-3,4-dehydro-L-proline benzyl ester **7** was added 50 mL THF, 5 mL H2O and 1.65 g (1.2 eq.; 29.4 mmol) KOH. The mixture was stirred for 2 h at rt. The mixture was concentrated by vacuum until dry in which 20 mL of water was added to dissolve the remaining residue. The pH of this mixture was adjusted to ~2 by addition of $H_3PO_4$ and NaCl was added until saturated. The contents were then extracted with EtOAc (3 x 200 mL). The ethyl acetate washes were combined and dried over $Na_2SO_4$, filtered and concentrated via rotovap to a yellow oil. The product, 6.05 g, N-Cbz-3,4-dehydro-L-proline **7a** was isolated by silica gel (60% hexanes, 39% ethyl acetate, 1% AcOH; $R_f$ = 0.2) a quantitative yield. The material was used in the next step without further purification.

*N-Cbz-trans-3,4-epoxy-L-proline* **8c**. To 2.37 g (9.60 mmol) of N-Cbz-3,4-dehydro-L-proline **7a** was added 20 mL $CH_2Cl_2$, 4.73 g (2 eq; 19.2 mmol) of 70% mCPBA, and 10 mg of (0.4% weight) 2,6-ditertbutyl-4-methyl phenol. The mixture was then refluxed under argon balloon overnight. Another 2.00 g (0.85 eq) of mCPBA was added to the reaction mixture and reflux was continued overnight. The mixture, now with white precipitate, was cooled over ice bath and filtered using 20 mL cold $CH_2Cl_2$ to wash. The product, 1.68 g of N-Cbz-*trans*-3,4-epoxy-L-proline **8c,** was separated from the crude mixture consisting of much reacted or unreacted mCPBA by a series of two silica gel columns; the first using 50% hexanes, 49% ethyl acetate, 1% AcOH as a mobile phase and the second using 96% $CH_2Cl_2$, 3% MeOH, 1% AcOH. Recovered epoxide product resulted in a yield of 70%, however, this represents only isolated epoxide from two columns with remaining epoxide eluting along with mCPBA; a higher yield is possible with further purification via flash chromatography and recrystallization. $^{1}$H-NMR (500 MHz, $CDCl_3$) (2 conformational isomers) δ 10.30 (b, 1H [COOH]), 7.35-7.26 (m, 5H), 5.18-5.11 (m, J = 20.8, 12.5 Hz, 1H), 5.12 (s, 1H), 4.75 and 4.68 (s, 1H), 3.97 and 3.92 (d, J = 12.7 Hz, 1H), 3.88 and 3.85 (d, J = 2.7 Hz, 1H), 3.69 (s, 1H), 3.56-3.53 (d, J = 12.7 Hz, 1H).

*(2S, 3S, 4R) N-Cbz-trans-3-hydroxy-cis-4-phenyl-*ʟ*-proline* ***9c*** *and (2S, 3S, 4R) N-Cbz-cis-3-phenyl-trans-4-hydroxy-*ʟ*-proline* ***9d***. To a flame dried round bottom flask was added 0.143 g (1.60 mmol; 3 eq) of CuCN (dried by high vacuum) and 5 mL dry ($MgSO_4$ treated) $Et_2O$. The flask was capped with a septum, purged air vacuum / argon balloon, and chilled to -40˚C (acetone bath / cold finger controled). Once cooled, 1.60 mL (3.19 mmol; 6 eq) of 2.0 M phenyl-lithium in THF was added and the mixture was then allowed to warm to -10˚C and kept at this temperature for ~15 min and then returned to -40˚C where 0.140 g (0.532 mmol) of N-Cbz-*trans*-3,4-epoxy-ʟ-proline **8c** in 2 mL dry ($MgSO_4$ treated) $Et_2O$ also at -40˚C was added via sterile syringe. The flask was then allowed to warm to -10˚C and stir overnight. After stirring overnight, the reaction mixture was allowed to warm to 0˚C and was quenched by the addition of 20 mL saturated $NH_4Cl$. The mixture was then warmed to rt, the pH was adjusted to ~2 by addition of $H_3PO_4$ and salted with NaCl until saturated. The products were then extracted with EtOAc (3 x 100 mL). The organic washes were combined, dried over $Na_2SO_4$, and concentrated to a solid residue via rotovap. The products, 0.171 g of (2S, 3S, 4R) N-Cbz-*trans*-3-hydroxy-*cis*-4-phenyl-ʟ-proline **9c** and (2S, 3S, 4R)-N-Cbz-*cis*-3-phenyl-*trans*-4-hydroxy-ʟ-proline **9d** were isolated as a regioisomer mixture (ratio ~2 : 3, as determined by $^{1}$H-NMR) by a series of silica gel columns (50% hexanes, 49% ethyl acetate, 1%

AcOH) for the first column followed by a second column ( 96% $CH_2Cl_2$, 3% MeOH, 1% AcOH), a 94% yield. The material was used in the next step without further purification.

*(2S, 3S, 4R) trans-3-hydroxy-cis-4-phenyl-L-proline* ***10c*** *(CSE104) and (2S, 3S, 4R) cis-3-phenyl-trans-4-hydroxy-L-proline* ***10d*** (CSE103). To a 0.200 g (0.586 mmol) regioisomer mixture of N-Cbz-*trans*-3-hydroxy-*cis*-4-phenyl-L-proline **9c** and N-Cbz-*cis*-3-phenyl-*trans*-4-hydroxy-L-proline **9d** was added 3 mL MeOH. The mixture was transferred to a pressure flask containing 20.0 mg (0.1 eq weight) 10% Pd / carbon, 0.5 mL $H_2O$ and 23.0 mg (0.5 eq: 0.298 mmol) $NH_4OAc$. The flask was placed on a Parr shaker, the air was removed by vacuum, and replaced with $H_2$ (g) to 40 psi. The apparatus was allowed to shake for 3 h at rt. The pressure flask was vented and the contents were diluted with 30 mL water and filtered through CM-cellulose using excess water for consecutive washes. The filtrate was then frozen and lyophilized until dry. The remaining residue was washed with EtOAc : $CH_2Cl_2$ 1:1 (3 x 20 mL), and dried leaving 0.108 mg (89% yield) of a off white solid, a crude mixture of *trans*-3-hydroxy-*cis*-4-phenyl-L-proline and *cis*-3-phenyl-*trans*-4-hydroxy-L-proline with ratios of 2 : 3 (as determined by $^1H$ NMR), respectively. A portion of this crude was separated by reverse phase HPLC (C18, 250 mm x 21.2 mm, 10μ) (92% 0.05 M $NH_4OAc$, 8% acetonitrile, 9 mL / min). Fractions containing a specific isomer were combined and concentrated by lyophilization. Once dry, 20 mL of water was added to dissolve the remaining residue and the solution was again frozen and lyophilized until dry (removes remaining $NH_4OAc$). This was repeated until $NH_4OAc$ was no longer present affording *trans*-3-hydroxy-*cis*-4-phenyl-L-proline **10c** (CSE104) and *cis*-3-phenyl-*trans*-4-hydroxy-L-proline **10d** (CSE103) as separate white solids.

*(2S, 3S, 4R) trans-3-hydroxy-cis-4-phenyl-L-proline* **10c** (CSE104). $^1$H-NMR (500 MHz, $D_2O$ 4.75) δ 7.41-7.38 (dd, J = 7.6, 7.1, 2H), 7.33-7.30 (m, J = 7.6 Hz, 3H), 4.63-4.61 (dd, J = 8.8, 4.4 Hz, 1H, H-2), 4.21-4.20 (d, J = 5.6 Hz, 2H, H-1), 3.91-3.86 (dd, J = 9.8, 5.4 Hz, 1H, H-5), 3.58-3.51 (m, 2H, H-3 & 4). $^{13}$C-NMR (500 MHz, $D_2O$) δ 171.0, 137.1, 129.1, 127.8, 127.0, 76.4, 64.6, 50.8, 47.9. HRMS *m/e* calcd. For $C_{11}H_{14}NO_3$ = 208.0974, found 208.0982. $[\alpha]^{22}_{Na}$ = -43° (c = 0.08, EtOH).

*(2S, 3S, 4R) cis-3-phenyl-trans-4-hydroxy-L-proline* ***10d*** (CSE103). $^1$H-NMR (500 MHz, $D_2O$ 4.75) δ 7.42-7.39 (dd, J = 7.3, 7.1 Hz, 2H), 7.35-7.31 (m, J = 7.6 Hz, 3H), 4.52-4.49 (dd, J = 12.7, 6.4 Hz, 1H, H-3), 4.21 (d, J = 8.8 Hz, 1H, H-1), 3.66-3.62 (dd, J = 11.5, 6.1 Hz, 1H, H-5), 3.38-3.31 (m, J = 8.8, 6.1 Hz, 2H, H-2 & 4). $^{13}$C-NMR (500 MHz, $D_2O$) δ 172.6, 136.8, 129.1, 127.9, 127.6, 75.4, 64.7, 55.8, 49.6. HRMS *m/e* calcd. For $C_{11}H_{14}NO_3$ = 208.0974, found 208.0996. $[\alpha]^{22}_{Na}$ = -16° (c = 0. 23, $H_2O$), $[\alpha]^{22}_{Na}$ = -15° (c = 0.24, EtOH).

*(2S, 3S, 4R) N-Cbz-trans-3-hydroxy-cis-4-methyl-L-proline* ***9a*** *and (2S, 3S, 4R) N-Cbz-cis-3-methyl-trans-4-hydroxy-L-proline* ***9b***. To a flame dried round bottom flask was added 0.730 g (8.50 mmol; 2.5 eq) of CuCN (dried by high vacuum) and 10 mL dry ($MgSO_4$ treated) $Et_2O$. The flask was capped with a septum, purged air with vacuum, filled with argon balloon, and cooled to -60˚C (acetone bath / cold finger controled). Once cooled, 10.0 mL (16.3 mmol; 5 eq) of 1.6 M methyl-lithium in $Et_2O$ was added via syringe and the mixture was allowed to stir for ~15 min until all the CuCN appeared to be dissolved (a clear colorless solution). Once equilibrated, 0.840 g (3.20 mmol) of N-Cbz-*trans*-3,4-epoxy-L-proline **8c** in 10 mL dry ($MgSO_4$ treated) $Et_2O$ at -60˚C was added via sterile syringe. The flask was then allowed to warm to -40˚C and stir overnight under argon balloon. After stirring overnight, the reaction mixture was allowed to warm to 0˚C and was

quenched by the addition of 30 mL saturated $NH_4Cl$. The mixture was then warmed to rt, the pH was adjusted to ~2 by addition of $H_3PO_4$ and salted with NaCl until saturated. The products were then extracted with of EtOAc (3 x 100 mL). The organic washes were then combined, dried over $NaSO_4$, and concentrated to a crude regioisomer mixture, 0.766 g. The products, N-Cbz-*trans*-3-hydroxy-*cis*-4-methyl-L-proline and N-Cbz-*cis*-3-methyl-*trans*-4-hydroxy-L-proline were isolated as a regioisomer mixture (ratios of 1:1 as determined by $^1H$ NMR) by silica gel from two columns (50% hexanes, 49% ethyl acetate, 1% AcOH and 96% $CH_2Cl_2$, 3% MeOH, 1% AcOH) an 86% yield and recovery of unreacted epoxide starting material. A portion of these regioisomers were separated by reverse phase HPLC (C18, 250 mm x 21.2 mm, 10 μ) (85% 0.05 M $NH_4OAc$, 15% acetonitrile, 9 mL/min) at a retention time of 22 min for (2S, 3S, 4R) N-Cbz-*cis*-3-methyl-*trans*-4-hydroxy-L-proline **9b** and 24 min for (2S, 3S, 4R) N-Cbz-*trans*-3-hydroxy-*cis*-4-methyl-L-proline **9a**. The fractions containing an individual isomer were combined and concentrated by lyophilization to afford the product as white solid. Each separate isomer was used in the next synthetic step without further purification.

*(2S, 3S, 4R) cis-3-methyl-trans-4-hydroxy-L-proline* ***10b*** (CSE109). To 70.0 mg (0.251 mmol) of N-Cbz-*cis*-3-methyl-*trans*-4-hydroxy-L-proline **9b** was added 2 mL MeOH.. The mixture was transferred to a pressure flask containing 7.0 mg (0.1 eq. weight) 10% Pd/carbon, and 6.3 (0.5 eq) $NH_4OAc$. The flask was placed on a Parr shaker, the air was removed by vacuum, and replaced by 40 psi $H_2$ (g). The apparatus was allowed to shake for ~2 h at rt. The pressure flask was vented and the contents were diluted with 30 mL water and filtered through CM-cellulose using excess water for consecutive washes. The filtrate was then frozen and lyophilized until dry. The remaining residue was washed with 1:1 EtOAc : $CH_2Cl_2$ ( 3 x 20 mL), resuspended in 10 mL water

and again dried by lyophilization leaving 33.0 mg of (2S, 3S, 4R) *cis*-3-methyl-*trans*-4-hydroxy-ʟ-proline **10b** (CSE109) as a white solid, a 92% yield. $^1$H-NMR (500 MHz, $D_2O$ 4.75) δ 4.09 (dd, J = 4.9, 2.4 Hz, 1H, H-3), 3.66 (d, J = 5.9 Hz, 1H, H-1), 3.50-3.46 (dd, J = 12.2, 4.9 Hz, 1H, H-5), 3.29-3.26 (dd, J = 12.2, 2.4 Hz, 1H, H-4), 2.37-2.33 (dd, J = 6.8, 5.9 Hz, 1H, H-2), 1.13 (d, J = 6.8 Hz, 3H, methyl). $^{13}$C-NMR (500 MHz, $D_2O$) δ 176.2, 77.4, 68.3, 52.8, 47.6, 18.6. HRMS *m/e* calcd. For $C_6H_{12}NO_3$ = 146.0817, found 146.0822. $[\alpha]^{22}_{Na}$ = -4° (c = 0.25, $H_2O$).

*(2S, 3S, 4R) trans-3-hydroxy-cis-4-methyl-ʟ-proline* ***10a*** (CSE110). To 80.0 mg (0.286 mmol) of N-Cbz-*trans*-3-hydroxy-*cis*-4-methyl-ʟ-proline **9a** was added 5 mL MeOH.. The mixture was transferred to a pressure flask containing 20.0 mg (0.25 weight %) 10% Pd/carbon. The flask was placed on a Parr shaker, the air was removed by vacuum, and replaced by 40 psi $H_2$ (g). The apparatus was allowed to shake for ~24 h at rt. The pressure flask was vented and the contents were diluted with 30 mL water and filtered through CM-cellulose using excess water for consecutive washes. The filtrate was then frozen and lyophilized until dry. The remaining residue was washed with 1:1 EtOAc / $CH_2Cl_2$ (3 x 20 mL), resuspended in 10 mL water and again dried by lyophilization leaving 34.0 mg of (2S, 3S, 4R) *trans*-3-hydroxy-*cis*-4-methyl-ʟ-proline **10a** (CSE110) as a white solid, a 82% yield. $^1$H-NMR (500 MHz, $D_2O$ 4.75) δ 4.27 (d, J = 4.4 Hz, 1H, H-2), 4.20 (d, J = 4.40 Hz, 1H, H-1), 3.65-3.61 (dd, J = 11.7, 6.60 Hz, 1H, H-4), 3.01-2.99 (dd, J = 11.7, 3.4 Hz, 1H, H-5), 2.42-2.39 (dd, 6.6 Hz, 3.4 Hz, 1H, H-3), 1.00 (d, J = 7.3 Hz, 3H, methyl). $^{13}$C-NMR (500 MHz, $D_2O$) ppm 170.7, 76.3, 65.0, 49.8, 40.6, 15.4. HRMS *m/e* calcd. For $C_6H_{12}NO_3$ = 146.0817, found 146.0829. $[\alpha]^{22}_{Na}$ = -12 ° (c = 0.31, $H_2O$).

*(2S, 3S, 4R) N-Cbz-trans-3-hydroxy-cis-4-phenoxy-*L*-proline* ***11a*** *and (2S, 3S, 4S) N-Cbz-cis-3-phenoxy-trans-4-hydroxy-*L*-proline* **11b**. To a round bottom flask containing 1.00 g (7.34 eq; 10.6 mmol) of phenol in 4 mL dry THF (distilled) was added 0.245 g (0.95 eq) of NaH slowly. The flask was then cooled over ice bath and argon balloon. To this mixture was added 0.381 g (1.45 mmol) N-Cbz-*trans*-3,4-epoxy-L-proline **8c** in 2 mL anhydrous THF (distilled) at 0˚C. The mixture was then allowed to warm to rt and then heated to reflux under condenser and argon balloon and stirred overnight. The reaction was then cooled on ice bath and 20 mL of 10% $H_3PO_4$ (aq) was added to quench the reaction. The mixture was then salted with NaCl until saturated and extracted with EtOAc (3 x 100 mL). The organic extracts were combined, dried over $NaSO_4$, and concentrated to a crude solid residue. The products, 0.330 g of N-Cbz-*trans*-3-hydroxy-*cis*-4-phenoxy-L-proline **11a** and N-Cbz-*cis*-3-phenoxy-*trans*-4-hydroxy-L-proline **11b** were isolated as a regioisomer mixture (ratios 12 : 1, respectively) by silica gel (50% hexanes, 49% ethyl acetate, 1% AcOH) and ( 96% $CH_2Cl_2$, 3% MeOH, 1% AcOH) from two columns to afford the product, a 64% yield not including recovered starting material. The regioisomer mixture was used in the next step without further purification.

*(2S, 3S, 4R) trans-3-hydroxy-cis-4-phenoxy-*L*-proline* ***12a*** (CSE118) *and (2S, 3S, 4S) cis-3-phenoxy-trans-4-hydroxy-*L*-proline* ***12b*** (CSE117). To a 0.550 g (1.54 mmol) regioisomer mixture of N-Cbz-*trans*-3-hydroxy-*cis*-4-phenoxy-L-proline **11a** and N-Cbz-*cis*-3-phenoxy-*trans*-4-hydroxy-L-proline **11b** was added 10 mL MeOH. The mixture was transferred to a pressure flask containing 83.0 mg (0.15 eq. weight) 10% Pd/carbon, 0.2 mL $H_2O$ and 60.0 mg (0.5 eq: 0.77 mmol) $NH_4OAc$. The flask was placed on a Parr shaker, the air was removed by vacuum, and replaced by 50 psi $H_2$ (g). The apparatus was allowed to shake overnight at rt. The pressure flask was vented and the

contents were diluted with 30 mL water and filtered through CM-cellulose using excess water for consecutive washes. The filtrate was then frozen and lyophilized until dry. The remaining residue was washed with 1:1 EtOAc / $CH_2Cl_2$ (3 x 20 mL), and dried leaving 0.308 mg of an off white solid, a crude mixture of (2S, 3S, 4R) *trans*-3-hydroxy-*cis*-4-phenoxy-L-proline and *cis*-3-phenoxy-*trans*-4-hydroxy-L-proline (12:1 respectively), a 90% yield. A portion of the product mixture was separated by reverse phase HPLC (C18, 250mm x 21.2mm, 10µ) (92% 0.05 M $NH_4OAc$, 8% acetonitrile, 9 mL/min). Collected fractions containing a single isomer were combined and concentrated by lyophilization. Once dry, 20 mL of water was added to redissolve the remaining residue and the solution was again frozen and lyophilized until dry (removes remaining $NH_4OAc$). This was repeated until $NH_4OAc$ was no longer present leaving (2S, 3S, 4R) *trans*-3-hydroxy-*cis*-4-phenoxy-L-proline **12a** (CSE118) or (2S, 3S, 4S) *cis*-3-phenoxy-*trans*-4-hydroxy-L-proline **12b** (CSE117) as a white solid.

*(2S, 3S, 4R) trans-3-hydroxy-cis-4-phenoxy-L-proline* **12a**. $^1$H-NMR (500 MHz, $D_2O$ 4.75) δ 7.33-7.30 (dd, J = 7.3, 7.3 Hz, 2H), 7.04-7.01 (dd, J = 8.3, 7.3 Hz, 1H), 6.93-6.92 (d, J = 8.3 Hz, 2H), 4.92 (d, J = 3.0 Hz, 1H, H-3), 4.67 (s, 1H, H-2), 4.10 (s, 1H, H-1), 3.76-3.73 (dd, J = 13.2, 3.0 Hz, 1H, H-4), 3.72-3.70 (d, J = 13.2 Hz, 1H, H-5). $^{13}$C-NMR (500 MHz, $D_2O$) δ 173.0, 157.8, 132.4, 125.0, 118.4, 81.9, 78.1, 70.3, 51.7. HRMS *m/e* calcd. For $C_{11}H_{14}NO_4$ = 224.0923, found 224.0937. $[\alpha]^{22}_{Na}$ = -81° (c = 0.1, EtOH).

*(2S, 3S, 4S) cis-3-phenoxy-trans-4-hydroxy-L-proline* **12b**. $^1$H-NMR (500 MHz, $D_2O$ 4.75) δ 7.33-7.30 (dd, J = 7.6, 7.6 Hz, 2H), 7.05-7.02 (dd, J = 8.1, 7.6 Hz, 1H), 7.00-6.98 (d, J = 8.1 Hz, 2H), 5.00 (d, J = 4.3 Hz, 1H, H-2), 4.59 (d, J = 4.3 Hz, 1H, H-1), 4.56 (d, J = 4.0 Hz, 1H, H-3), 3.73-3.69 (dd, J =

13.0, 4.0 Hz, 1H, H-5), 3.36 (d, J = 13.0 Hz, 1H, H-4). $^{13}$C-NMR (500 MHz, $D_2O$) δ 169.6, 156.1, 129.8, 122.5, 115.9, 81.2, 71.8, 64.2, 51.0. HRMS *m/e* calcd. For $C_{11}H_{14}NO_4$ = 224.0923, found 223.0911. $[\alpha]^{22}_{Na}$ = -15° (c = 0.08, EtOH).

*N-Trityl-4-keto-L-proline ethyl ester* **24a**. To a 100 mL round bottom flask containing 20 mL $CH_2Cl_2$ was added 0.54 mL (6.20 mmol; 1.2 eq) of $COCl_2$ in 2 mL $CH_2Cl_2$. The flask was then chilled to -60˚C (acetone bath and cold finger) under argon balloon. To this was added 0.88 mL dry (molecular civ treated) DMSO in 1 mL dry ($MgSO_4$ treated) $CH_2Cl_2$ drop wise. The mixture was allowed to stir for 15 min before being brought up to -30˚C where 2.02 g (6.17 mmol) of N-trityl-*trans*-4-hydroxy-L-proline ethyl ester **3a** in 4 mL $CH_2Cl_2$ was added dropwise over 10 min. The resulting solution was allowed to stir for 1 h at -30˚C under argon balloon. To complete the reaction, 3.6 mL (> 5 eq) of triethylamine was slowly added. The flask was then allowed to warm to rt where 100 mL of water was added. The mixture was then extracted with $Et_2O$ (3 x 100 mL). The organic extracts were combined, dried with $Na_2SO_4$ and concentrated via rotovap to a solid crude. The remaining crude was then immediately separated by silica gel (90% hexanes, 9% ethylacetate, 1% TEA, $R_f$ = 0.2). The fractions containing the product were combined, concentrated, and placed on a high vacuum leaving 1.51 g of N-trityl-4-keto-L-proline ethyl ester **24a**, as a white foam, a yield of 75%. This product, once isolated, was then used within 4-6 h for the addition step (trityl-keto-proline intermediate readily decomposes). $^1$H-NMR (400 MHz, $CDCl_3$ at 7.27 ppm) δ 7.58-7.56 (d, J = 8.1 Hz, 6H), 7.31-719 (m, 9H), 4.30-4.18 (m, 3H), 3.86-3.81 (d, J = 19.8 Hz, 1H), 3.54-3.49 (d, J = 19.8 Hz, 1H), 1.88-1.84 (d, J = 18.3 Hz, 1H), 1.34-1.31 (dd, J = 13.9, 6.6 Hz, 3H), 1.12-1.05 (dd, J = 18.3, 10.3, 1.5 Hz, 1H). $^{13}$C-NMR (400 MHz, $CDCl_3$ at 77.00 ppm) δ 215.7, 173.5, 143.6, 128.88, 128.2, 127.9, 126.8, 77.9, 61.1, 60.1, 57.1, 40.2, 14.2.

*N-Trityl-cis-4-methyl-trans-4-hydroxy-*L*-proline ethyl ester* **25a**. To a flame dried / argon purged flask containing 0.123 g (5.00 mmol; 2 eq) of Mg metal in 2 mL diethyl ether was added 0.315 mL (5.00 mmol; 2 eq) of iodomethane over 10 min. The mixture was allowed to stir for ~ 20 min until when it was observed that most or all of the Mg metal was consumed. The flask was cooled to -20˚C under argon balloon where it was then transferred to a flask containing 0.180 mg (1.27 mmol; 0.5 eq) of CuI (dried by vacuum) at -20˚C. The mixture was allowed to stir for ~15 min. after which 1.00 g (~2.50 mmol) of N-trityl-4-keto-L-proline ethyl ester **24a** in 6 mL dry $Et_2O$ was added and allowed to stir overnight. The reaction was then quenched by the addition of 15 mL saturated $NH_4Cl$ solution, and diluted with 100 mL $Et_2O$. The organic phase was washed with water (3 x 100 mL), dried over $Na_2SO_4$ and concentrated via rotovap to a light brown foam. Immediately following the addition step, the product, 0.624 g N-Trityl-*cis*-4-methyl-*trans*-4-hydroxy-L-proline ethyl ester **25a**, was isolated by silica gel (95% hexanes, 4% ethyl acetate, 1% TEA; $R_f$ = 0.12), a yield of 60%. $^1$H-NMR (400 MHz, $CDCl_3$ at 7.27 ppm ) δ 7.61-7.60 (d, J = 7.3 Hz, 6H), 7.32-728 (dd, J = 8.1, 7.3 Hz, 6H), 7.21-7.18 (dd, J = 7.3 Hz, 3H), 4.03-3.90 (m, 3H ), 3.63-3.60 (d, J = 12.4 Hz, 1H), 2.99-2.96 (d, J = 12.4 Hz, 1H), 1.93-1.88 (dd, J = 13.2, 6.6 Hz, 1H), 1.71-1.31 (dd, J = 13.2, 8.1 Hz, 1H), 1.25 (s, 3H), 1.15-1.12 (dd, J = 7.3, 7.3 Hz, 3H). $^{13}$C-NMR (400 MHz, $CDCl_3$ at 77.00 ppm) δ 175.9, 144.0, 129.4, 127.8, 126.5, 77.8, 77.3, 63.7, 62.1, 60.4, 46.7, 26.2, 14.0.

*cis-4-methyl-trans-4-hydroxy-*L*-proline* **26** (CSE95). To 0.111 g (0.276 mmol) of N-Trityl-*cis*-4-methyl-*trans*-4-hydroxy-L-proline ethyl ester **25a** was added 2 mL of THF, 0.2 mL $H_2O$ and 20.0 mg (90%, 1.2 eq) KOH. The mixture was allowed to stir for 2 h at rt. The solution was then concentrated to a solid residue by vacuum, and washed with water (2 x 10 mL). The remaining residue was resuspended in 2 mL THF and 0.33 mL (2.5 eq) of 2 M HCl was added. After 1 h of stirring the solution was diluted with 10 mL water. The THF was removed by rotovap leaving mostly water and dissolved product. The sample was frozen and lyophilized to concentrate. The remaining residue was washed 1:1 EtOAc / $CH_2Cl_2$ (3 x 10 mL). The sample was then diluted with 5 mL water and isolated by cation exchange resin (sulfonic acid resin) using 10% $NH_4OAc$ to elute the final product, *cis*-4-methyl-*trans*-4-hydroxy-L-proline **26** (CSE95). The collected fractions were combined and concentrated by lyophilization to afford 32 mg of the product, an 83% yield. $^1$H-NMR (500 MHz, $D_2O$ at 4.75) δ 4.30-4.25 (dd, J = 10.4, 7.8 Hz, 1H, H-1), 3.22 (s, 2H, H-4 & 5), 2.40-2.34 (dd, J = 13.6, 8.4 Hz, 1H, H-2), 1.98-1.92 (dd, J = 13.6, 10.4 Hz, 1H, H-3), 1.38 (s, 3H). $^{13}$C-NMR (500 MHz, $D_2O$) δ 175.1, 78.0, 61.0, 56.8, 42.9, 22.9. HRMS *m/e* calcd. For $C_6H_{12}NO_3$ = 146.0817, found 146.0803. $[\alpha]^{22}_{Na}$ = -60° (c = 0.35, MeOH).

*N-Cbz-4-keto-L-proline* **24b**. To a round bottom flask containing 14.0 g (52.8 mmol) of N-Cbz-*trans*-4-hydroxy-L-proline **3** was added 26 mL acetone. The solution was cooled on ice bath where 8.12 mL (0.106 mmol; 2 eq) of Jones reagent ($CrO_3$, $H_2SO_4$, 1.3 M in water) was added. The mixture was allowed to warm to rt and stir for 3 h where the reaction appeared to be complete, as determined by TLC. The reaction was quenched by the addition of 6 mL isopropanol. After stirring for another 2 h the flask was placed on a glass socket rotovap and the isopropanol/acetone was removed by reduced pressure. The remaining green residue was then diluted into 200 mL water, salted with NaCl and extracted with EtOAc (3 x 200 mL). The organic extracts were combined, dried with $MgSO_4$, filtered and concentrated to an oil. The remaining yellow oil was then separated by silica gel (90% $CH_2Cl_2$, 9% MeOH, 1% AcOH, $R_f$ = 0.3). The fractions containing the product were combined and concentrated by rotovap to a solid residue and then recrystallized from $CH_2Cl_2$ leaving 9.86 g (37.5 mmol) of N-Cbz-4-keto-L-proline **24b**, a

71% yield. $^{1}H$ and $^{13}C$ NMR spectra were in agreement with spectra previously reported [Bridges *et al* 1991].

*N-Cbz-trans-4-methyl-cis-4-hydroxy-L-proline* **25b**. To a flame dried / argon purged flask containing 0.231 g (9.40 mmol; 2.5 eq) Mg metal in 2 mL dry diethyl ether was added 0.530 mL (8.40 mmol; 2.2 eq) of iodomethane slowly. The mixture was allowed to stir for ~ 20 min until when it was observed that most or all of the Mg metal was consumed. The mixture was then cooled to -20˚C under argon balloon. To this was added 0.5 eq of CuI and allowed to stir for 15 min. To this prepared Gilman reagent was added 1.00 g (2.50 mmol) of N-Cbz-4-keto-L-proline **24b** in 5 mL cold dry $Et_2O$. The mixture was allowed to return to rt and heated at 35 C for 2 h. The reaction was then quenched by the addition of 15 mL saturated $NH_4Cl$ aq and allowed to stir an additional 30 min. The mixture was then acidified with HCl to pH ~2, salted with NaCl and extracted with EtOAc (3 x 100 mL). The organic extracts were then combined, dried with $MgSO_4$, filtered, and concentrated to a light brown residue. The product, 0.440 g (1.58 mmol) of N-Cbz-*trans*-4-methyl-*cis*-4-hydroxy-L-proline **25b**, was isolated by a silica gl (85% $CH_2Cl_2$, 14% MeOH, 1% AcOH; $R_f$ = 0.25) with a yield of 41%; not including full recovery of unreacted starting material. The isolated product was then used in the next deprotection step without further purification. $^{1}H$-NMR (400 MHz, CDCl3 at 7.27 ppm) δ 8.73 (b, 1H), 7.33-7.27 (m, 5H), 5.18-5.07 (m, 2H), 4.48-4.42 (dd, J = 15.4, 8.8 Hz, 1H), 3.70-3.64 (dd, J = 11.7 Hz, 1H), 3.37-3.32 (dd, J = 10.3 Hz, 1H), 2.25-2.15 (m, 2H), 1.37-1.35 (d, J = 8.1 Hz, 3H). $^{13}C$-NMR (400 MHz, $CDCl_3$ at 77.00 ppm) (2 conformational isomers) δ 176.8 and 176.6, 155.4 and 154.8, 135.9, 128.4 and 128.3, 128.0, 127.8 and 127.6, 76.8 and 76.2, 67.5 and 67.4, 59.4 and 59.3, 58.8 and 58.5, 43.4 and 42.4, 24.2 and 20.6.

*cis-4-hydroxy-trans-4-methyl-L-proline* **27** (CSE96). To 0.200 g (0.716 mmol) of N-Cbz-*cis*-4-hydroxy-*trans*-4-methyl-L-proline **25b** was added 2 mL MeOH which was transferred to a pressure flask containing 50 mg 10% Pd/carbon and 2 mL $H_2O$ and 0.2 eq $NH_4OAc$. This was then connected to a Parr shaker and the air was removed by vacuum. The container was filled with 50 psi $H_2$ (g) and the apparatus was allowed to shake overnight. The flask was then vented and the contents were diluted with 20 mL water and filtered through CM-cellulose. The filtrate was then frozen and concentrated by lyophilization to a light brown residue that was then washed with 1:1 EtOAc : $CH_2Cl_2$ (3 x 10 mL). The sample was then diluted with 5 mL water and isolated by cation exchange resin (sulfonic acid resin) using 10% ammonium acetate to elute the final product. The collected fractions were combined and concentrated by lyophilization to afford 62 mg of *cis*-4-hydroxy-*trans*-4-methyl-L-proline **27** (CSE96), a 60% yield. $^{1}$H-NMR (400 MHz, $D_2O$ 4.75) δ 4.55-4.52 (dd, J = 9.1, 3.9 Hz, 1H, H-1), 3.37-3.34 (d, J = 11.6 Hz, 1H, H-5), 3.20-3.17 (d, J = 12.3, 1H, H-4), 2.35 (m, H-2 and 3), 1.36 (s, 3H). $^{13}$C-NMR (500 MHz, $D_2O$) δ 76.8, 59.4, 57.1, 41.9, 22.4. HRMS *m/e* calcd. For $C_6H_{12}NO_3$ = 146.0817, found 146.0823. $[\alpha]^{22}_{Na}$ = -44° (c = 0.39, MeOH).
